# Analysis of a diverse wheat germplasm panel reveals a highly diverse introgression landscape and provides evidence for inter-homoeolog chromosomal recombination

**DOI:** 10.1101/2024.01.10.574987

**Authors:** Matthias Heuberger, Zoe Bernasconi, Esther Jung, Gerhard Herren, Victoria Widrig, Beat Keller, Javier Sánchez-Martín, Thomas Wicker

## Abstract

Agriculturally important genes are often introgressed into crops from closely related donor species or landraces. The gene pool of hexaploid bread wheat (*Triticum aestivum)* is known to contain numerous such “alien” introgressions. Recently established high-quality reference genome sequences allow prediction of the size, frequency, and identity of introgressed chromosome regions. Here, we characterise chromosomal introgressions in bread wheat using exome capture data from the WHEALBI collection. We identified 26,664 putative introgression segments of at least 2 Mb across 434 wheat accessions. Detailed study of the most frequent introgressions identified *T. timophevii* or its close relatives as a frequent donor species. Importantly, 118 introgressions of at least 10 Mb were exclusive to single wheat accessions, revealing that large populations need to be studied to assess the total diversity of the wheat pangenome. In one case, a 14 Mb introgression in chromosome 7D, exclusive to cultivar Pamukale, was shown by QTL mapping to harbor a recessive powdery mildew resistance gene. We identified multiple events where distal chromosomal segments of one subgenome were duplicated in the genome and replaced the homoeologous segment in another subgenome. We propose that these examples are the results of inter-homoeolog recombination. Our study produced an extensive catalogue of the wheat introgression landscape, providing a resource for wheat breeding. Of note, the finding that the wheat gene pool contains numerous rare, but potentially important introgressions and chromosomal rearrangements has implications for future breeding.

**Key message:** This study highlights the potential of rare introgressions, as demonstrated by a major QTL for powdery mildew resistance on chromosome 7D. It further shows evidence for inter-homoeolog recombination in wheat.

## Introduction

Bread wheat (*Triticum aestivum*) is a hexaploid species (genome formula AABBDD) that resulted from two allopolyploidization events. The A, B and D subgenomes each have seven chromosomes referred to as homoeologs (Glover et al. 2016). The first polyploidization occurred about 0.5 million years ago when *T. urartu* (AA) and a yet unknown donor, closely related to *Aegilops speltoides,* hybridized to form *T. turgidum* (AABB) (Marcussen et al. 2014). This tetraploid wheat, which began to be cultivated approximately 10,000 years ago in the Fertile Crescent, subsequently hybridised with *Ae. tauschii* to form bread wheat. Both these allopolyploidization events acted as genetic bottlenecks so bread wheat contains only a fraction of the genetic diversity found in the genomes of the three progenitor species. This reduced diversity is most pronounced in the D subgenome (Gaurav et al. 2021).

It is well known that wild relatives of crops and landraces are a rich resource of novel genes for exploitation in contemporary agriculture (Hajjar and Hodgkin 2007). The genetic diversity of this original gene pool was only partially incorporated into modern wheat varieties. One method to introduce genetic material from wild crop relatives is introgression breeding (Hao et al. 2020). Here, a crop variety is hybridised with a relative containing a desired trait. Genetic recombination between non-homologous chromosomes of wheat is suppressed mainly by the *Ph1* locus (Griffiths et al. 2006). Therefore, the *Ph1* locus is either removed before introgression breeding, or the cross is made with a species not containing the *Ph1* locus, such as *Ambylopyrum muticum* (King et al. 2017, 2019; Coombes et al. 2023). After a successful cross, the resulting offspring is subsequently backcrossed while selecting the offspring for the trait of interest. This is done to reduce potentially unwanted phenotypes originating from the donor genotype.

In bread wheat, introgression breeding has been widely used for many traits and genes of agronomic importance. Breeding for disease resistance has relied heavily on introgressions from wild or crop relatives. For example, the 1RS.1BL translocation where the short arm of wheat chromosome 1B is replaced by the short arm of rye (*Secale cereale*) chromosome 1R, contains genes conferring resistance to four different fungal pathogens: *Lr26* for resistance to leaf rust (*Puccinia triticina*), *Yr9* for resistance to yellow rust (*P. striiformis*), *Sr31* for resistance to stem rust (*P. graminis*), and *Pm8* for resistance to powdery mildew (*Blumeria graminis* f. sp*. tritici,* or *Bgt*) (Crespo-Herrera et al. 2017). Examples of other introgressed disease resistance genes are *Sr43* from *Thinopyrum elongatum* (Knott et al. 1977; Yu et al. 2023), *Lr9* (Wang et al. 2023), *Lr76*, *Yr70* (Bansal et al. 2020) from *Ae. umbellulata* and *Pm4b* from *T. carthlicum* (Sánchez-Martín et al. 2021).

Cytological analyses such as C-banding have been important tools to detect introgressions in specific genotypes and they rely on differential staining of heterochromatin (Badaeva et al. 1991; Orlovskaya et al. 2020). A molecular method to determine the presence of introgressions is simple sequence repeat (SSR) analysis, where the differential amplification of SSR markers informs about the presence of alien chromatin (Orlovskaya et al. 2020). The advent of next generation sequencing (NGS) and the availability of reference wheat genomes now enable whole genome analysis (Walkowiak et al. 2020). Genomic methods to identify introgressions include transposable element population analysis (Walkowiak et al. 2020; Wicker et al. 2022), k-mer-based approaches (https://github.com/Uauy-Lab/IBSpy, Ahmed et al. 2023) and single nucleotide polymorphism (SNP)-based methods (Cheng et al. 2019). Additionally, a straightforward way of introgression detection is the analysis of sequencing coverage depth (Keilwagen et al. 2019; Walkowiak et al. 2020; Keilwagen et al. 2022; Kale et al. 2022; Keilwagen et al. 2023). Here, genomic sequence reads (typically Illumina short reads) from a given wheat accession are mapped to a reference genome. In regions containing divergent haplotypes or introgressions, fewer sequence reads will map due to lower sequence homology. The method does not directly allow the differentiation between introgressions which are present in the reference genome and introgressions that are present in the sample. However, it is possible to overcome this limitation by using multiple genomes as reference.

To assess the current diversity of the modern wheat gene pool, several collections of wheat genotypes were assembled and made publicly available. In this study, we used the WHEALBI resource (Pont et al. 2019, www.whealbi.eu), a collection of 434 hexaploid genotypes which were categorised in the WHEALBI project into historic groups as landraces (deployment before 1935, N=95), old cultivars (registered from 1936 to 1965, N=99), cultivars (registered from 1966 to 1985, N=130) and current varieties (registered after 1986, N=99, 11 accessions are not categorised). By mapping publicly available exome capture reads from the WHEALBI collection onto the wheat reference genome (cv. Chinese Spring v2.1, Zhu et al. 2021), we identified a total of 26,664 putative introgressions across 434 wheat accessions. We scrutinised the most frequently occurring introgressions and identified potential donors. We also identified a large number of introgressions that are private to single genotypes. The practical value of the presented data sets was exemplified by QTL mapping of a recessive powdery mildew resistance gene on chromosome 7D embedded in a 14 Mb introgression that is only present in the Turkish cultivar Pamukale. Finally, we identified several duplicated segments in distal chromosomal regions, which we propose were the results of inter-homoeolog recombination. Our findings suggest that recombination between homoeologous chromosomes was a frequent event during wheat evolution.

## Materials and methods

### Mapping of exome capture and other reads

Genomes were first indexed with bwa index (v.0.7.17,Li 2013) using the -c flag. The exome capture reads were then mapped on the CS genome (IWGSC RefSeq v2.1) using bwa mem (v.0.7.17). The resulting sam file was then converted to .bam format and duplicate reads were removed with samtools (v.1.6, Danecek et al. 2021) using the functions sort fixmate and markup. Coverage ratio was then calculated using the deeptools (v.3.5.1, Ramírez et al. 2016) command bamCompare, using the mapping of CS exome capture reads as normalisation control with the flags: --minMappingQuality 30, --operation ratio, --binSize 500000. For GBS and WGS of wild relatives, the same method was used but with a bin size of 2 Mb.

### Detection of introgressions

Introgressions were automatically detected by a custom Perl script (https://github.com/matthias-heuberger/heubiSOFT/blob/main/assign_introgressions_WW), which evaluated for each bin if the coverage was below 0.5. Adjacent bins below this threshold were connected to form one putative introgression region. Note that this means if two regions of low coverage are separated by a region with coverage ratio above 0.5, these introgressions are counted separately. They might however not be truly independent introgression events.

### Determination of uniqueness of introgressions

To distinguish whether introgressions are unique or shared between accessions, they were classified based on whether their start and end points were the same in multiple accessions. Unique introgressions of size 10 Mb or more were further scrutinised manually. To be counted as unique either the start or the endpoint had to be at least 2 Mb from the start or endpoint of another introgression, or the coverage ratio across the entire introgression had to deviate at least +-30% from other introgressions in the same region.

### Determination of gene coverage

To determine the coverage of genes in the exome capture sequencing data mapping coverage of Chinese Spring against itself was analysed using the program featureCounts (v.2.0.0, Liao et al. 2014). The average coverage across the entire gene was then calculated using the read counts per gene obtained by featureCounts and the gene and read length. A gene was counted as sufficiently covered, if the gene had an average coverage of >=3 reads.

### Fungal material and infection tests

*Blumeria graminis* f. sp. *tritici* (*Bgt*) isolates Bgt_07004 and Bgt_10001 were used to evaluate Pamukale resistance. Both *Bgt* isolates were avirulent on Pamukale (no fungal growth) but virulent on the susceptible cultivars Frisal and Kanzler (leaves fully covered in fungal mycelia; Fig. 3b; at least 3 biological replicates were tested per *Bgt* isolate). To assess the resistance response, 3 cm leaf fragments were placed on water agar containing 500 ppm Benzimidazole and spray-inoculated with *Bgt*. The infection rate was evaluated 7-9 days after infection and described as the percentage of leaf area covered by fungal mycelia (ranging from 0 being completely resistant to 100 being completely susceptible). Other *Bgt* isolates used for phenotyping have been selected from a worldwide collection described in (Sotiropoulos et al. 2022).

### SNP based genotyping and filtering

Genomic DNA was extracted from leaf tissues of Frisal, Pamukale and the 133 F2 individuals resulting from their cross, using the CTAB method (Stein et al. 2001). Genotyping has been performed by SGS INSTITUT FRESENIUS GmbH TraitGenetics (https://sgs-institut-fresenius.de) with the Wheat Illumina Infinium 25K SNP Array. The SNP sequences were blasted against the genomes of 10 wheat cultivars (Walkowiak et al. 2020) and only the SNPs consistently mapping to the same chromosome for all tested wheat genomes were kept for further analyses. From the initial 24,146 SNP markers, 12,936 SNP markers passed this filtering step.

### Linkage map construction and QTL mapping

Further filtering and linkage map construction was done with the software QTL IciMapping, (v.4.2; https://isbreedingen.caas.cn/; Meng et al. 2015). First, SNP markers which were non-polymorphic or missing in the parents were removed by deletion using the “SNP” functionality. Redundant markers and markers with more than 50% missing rate were removed with the “BIN” functionality using the following parameters: p-value: 0.01, missing values considered. In the “MAP” functionality, chromosome and physical positions have been considered for linkage map construction. The final linkage map consists of 2,058 SNP markers and is available in Supp. Table S5. QTL mapping was performed with the R package R/qtl (v1.6), as described by (Müller et al. 2022). Briefly, the function read.cross() was used to import map position, genotypic (Supp. Table S5) and phenotypic data (Supp. Table S4) of the Pamukale x Frisal cross. Subsequently, jittermap() and calc.genoprob() commands were used to filter the genotypic data before performing QTL mapping with scanone(, model=”np”). The significance threshold was estimated with 500 permutations.

### Statistical analyses

Chi-square goodness of fit test has been performed with base R. Plots and other figures have been produced with R/RStudio (v.4.3.0 and 2023.03.1 respectively), using the package ggplot2 or base R.

### PCR amplifications across candidate breakpoints

The region +-3 kb of the suspected breakpoint of both homoeologous regions was extracted from IWGSC RefSeq v2.1. The two homoeologous regions were then aligned using the Smith-Waterman algorithm (Smith and Waterman 1981). Regions of the alignment with homoeolog specific SNPs or InDels were chosen as targets for primer design, which was carried out using Primer-BLAST (Ye et al. 2012) based on Primer3 (Untergasser et al. 2012). PCR amplifications were done using 50 ng of template DNA, using a homemade Pfux7 polymerase (Nørholm 2010) with the Phusion Green HC Buffer (F538L, Thermo Fisher). Primer sequences are listed in Supp. Table S7.

## Results

### A catalogue of wheat chromosomal introgressions

We used exome capture data from the 434 wheat accessions from the WHEALBI collection as a basis to identify introgressions. The exome capture data covers ∼1% of the genome and ∼59.8% of all annotated genes (Supp. Table S1). To identify candidate introgression regions, exome capture sequence reads from each wheat accession were mapped against IWGSC RefSeq v2.1 (cv. Chinese Spring, hereafter referred to simply as “Chinese Spring” (CS)). The resulting read coverage was normalised against the sequence coverage of reads from CS (hereafter referred to as “coverage ratio”). In regions where a sample and the reference have the same haplotype a coverage ratio of ∼1 was expected.

We defined introgressions as regions showing a coverage ratio smaller than 0.5 (i.e. the coverage was only one-half or less than CS in the same bin). Adjacent chromosome bins which were classified as introgressions were merged for determining the total number of introgressed segments. With this method introgressions were defined by comparison with CS and it was not possible to determine whether the introgression was present in the specific accession or in CS. One example of an introgression in Chinese Spring has been described on chromosome 6A and it originates from *T. monococcum* (Ahmed et al. 2023).

Using the coverage ratio cutoff of <0.5 we identified 26,644 putative introgressions of at least 2 Mb in size in the WHEALBI collection (i.e. four adjacent genomic bins, see methods). Introgressed regions span, on average, 4.12% of the genome of a given accession (median = 3.96%). The D subgenome has the smallest introgressed fraction with an average of 1.80% and a median of 1.76% (Fig. 1a). The median number of introgressions per accession is 60 (mean = 61.58), eight are larger than 5 Mb (mean = 9.21) and three are larger than 10 Mb (mean = 2.92). Strikingly, 400 of 433 accessions have five or more introgressions larger than 5 Mb and more than half (52.66%) of all accessions have at least three introgressions larger than 10 Mb. Finally, 2991 introgressions are private to single accessions, of which 902 are at least 5 Mb in size and 118 are at least 10 Mb (Supp. Table S2). Overall, these data indicate that multiple and large introgressions are quite common in members of the WHEALBI collection.

**Fig. 1.**
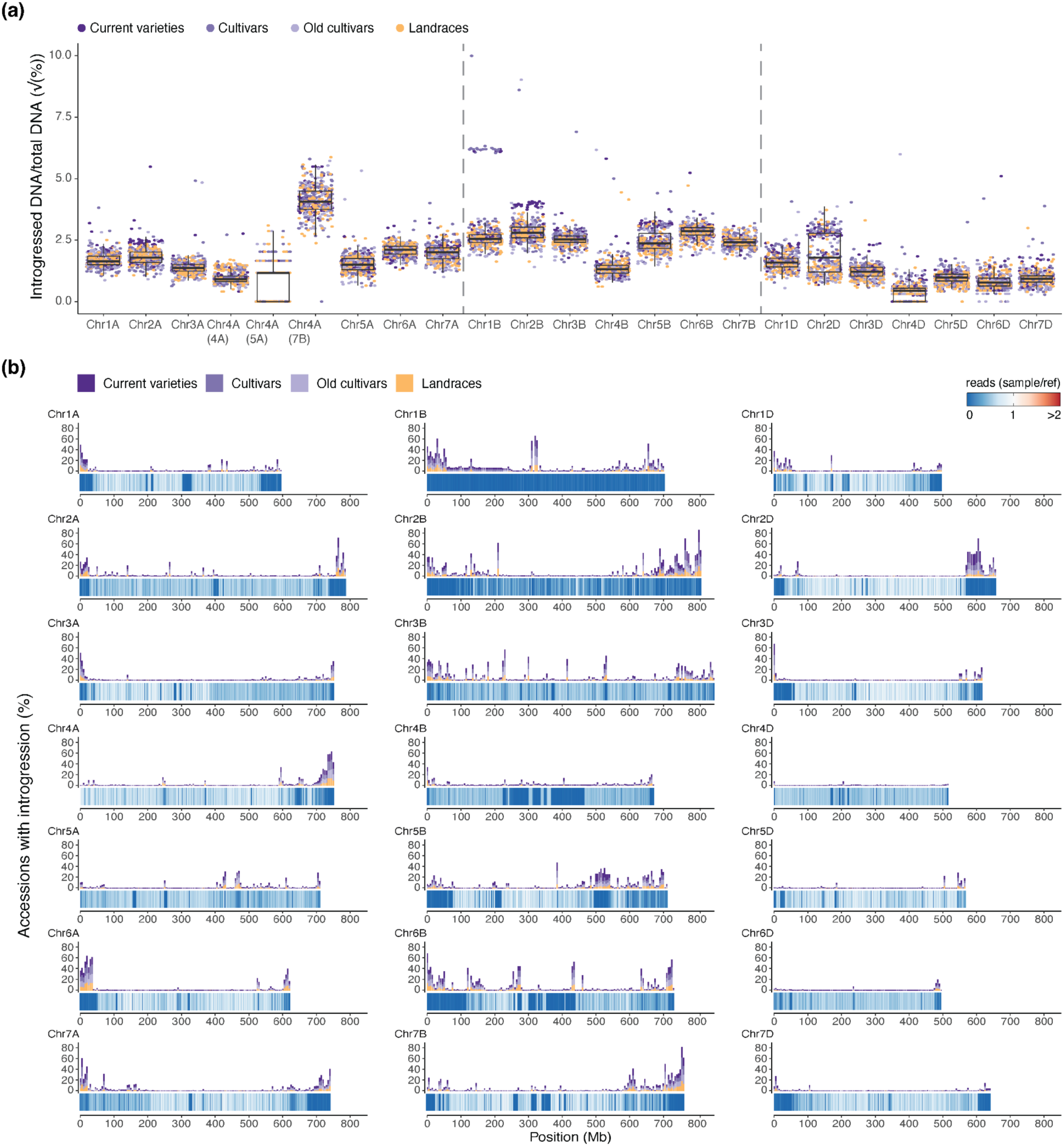
Overview of chromosomal introgressions in the WHEALBI collection. **(a)** Boxplot showing the percentage of each chromosome that is part of an introgression in hexaploid genotypes of the WHEALBI collection. Each dot corresponds to one wheat accession. Color indicates the group. Chromosome 4A was split into three parts, according to the subgenome ancestry of the different segments of chromosome 4A (Dvorak et al. 2018). Boxes indicate the inter-quartile range (IQR) with the central line indicating the median and the whiskers indicating the minimum and maximum values without outliers extending beyond - 1.5*IQR and maximum + 1.5*IQR. **(b)** Introgression frequencies along wheat chromosomes. For each chromosome the heatmap at the bottom shows the minimal sequence coverage in the respective 500 kb bin. The barchart on top shows the percentage of accessions having an introgression in this bin.

### Large chromosomal introgressions are frequent in the WHEALBI collection

We identified several introgressions from known donor species previously described in the literature (Fig. 2a-b). The largest introgression was in Riebesel St. 47-51 (WW-098), where the entire 700 Mb chromosome 1B is replaced by *S. cereale* (rye) chromosome 1R (727 Mb) (Friebe et al. 1987). Among the introgressions larger than 10 Mb, the most frequently observed is located on chromosome 2D from ∼573 Mb to ∼622 Mb (Fig. 1b, Fig. 2a, Supp. Fig. 1a). This introgression has been identified in multiple reference quality genome assemblies (RQAs) and has been suggested to originate from *A. markgrafii* (Walkowiak et al. 2020; Keilwagen et al. 2022; Supp. Fig. 1a). Strikingly, this introgression is found in the majority of current varieties and cultivars (87.9% and 52.3% respectively), but only in 12.6% of landraces. This introgression was previously reported to be enriched in elite genotypes and positively affects total plant biomass and number of grains per m^2^ under optimal conditions (Voss-Fels et al. 2019; Keilwagen et al. 2022). It was also shown to contain a homologue of a gene conferring resistance to yellow rust (Keilwagen et al. 2022).

**Fig. 2.**
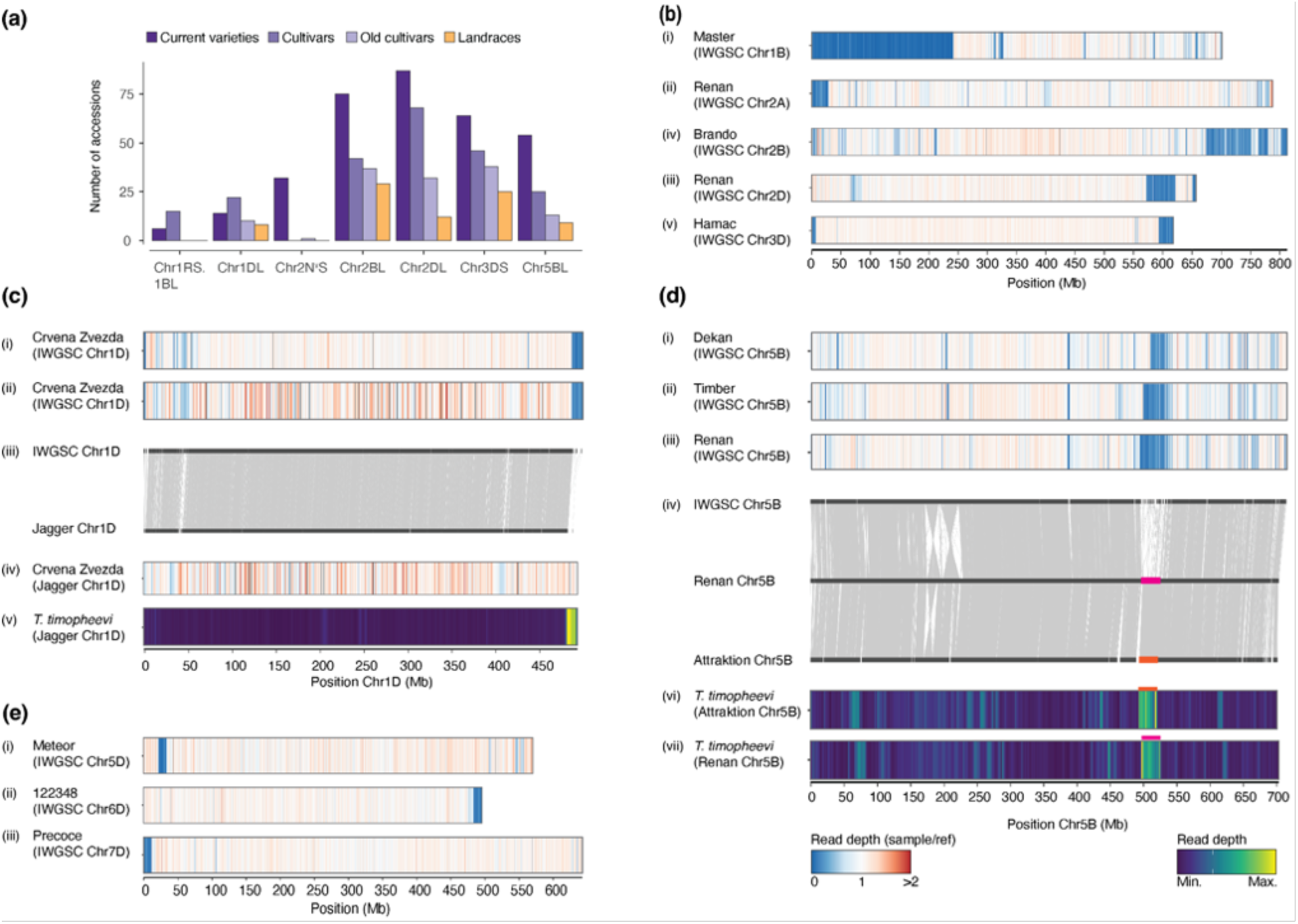
Detailed characterization of chromosomal introgressions **(a)** Barplot showing the number of accessions carrying introgressions. This includes the Chr.1RS.1BL translocation, the *Ae. ventricosa* introgression on Chr. 2A, the *T. timopheevii* introgression on Chr. 2B, the *Ae. markgrafii* introgression on Chr. 2D, the *Ae. comosa* /*Ae. uniarista* introgression on Chr. 3D (Keilwagen et al. 2022) and the two herein described introgression likely coming from *T. timopheevii* on Chr. 1D and Chr. 5B. **(b)** Introgression regions quantified in (a). Coverage of normalised exome capture reads of: (i) Master on Chr. 1B; (ii) Renan on Chr. 2A; (iii) Brando on Chr. 2B; (iv) Renan on Chr. 2D; and (v). Hamac on Chr. 3D. Values were calculated in 500 kb bins. **(c)** Candidate introgression region on Chr. 1D; (i) Coverage of normalised exome capture reads of Crvena Zvezda on Chr. 1D; (ii) coverage of normalised GBS reads of Crvena Zvezda on Chr. 1D; (iii) chromosomal synteny between CS Chr. 1D and Jagger Chr. 1D; (iv) coverage of normalised GBS reads of Crvena Zvezda on Jagger Chr. 1D; (v) *T. timopheevii* WGS reads mapped on Jagger Chr. 1D. Values for exome capture were calculated in 500 kb bins, GBS and WGS in 2 Mb bins **(d)** Candidate introgression region on chromosome 5B. Normalized coverage of: (i) Dekan exome capture reads; (ii) Timber exome capture reads and (iii) Renan exome capture reads mapped on CS Chr. 5B. (iv) Chromosomal collinearity between Chinese Spring, Renan and Attraktion chromosome 5B. *T. timopheevii* WGS reads mapped on chromosome 5B in Attraktion (v) and Renan 5B (vi). Values for exome capture were calculated in 500 kb bins and WGS in 2 Mb bins.

Another large introgression with a known donor species is found only in current varieties (32.3%), except for one old cultivar (Taferstat, WW-420). It is located in a ∼29 Mb segment at the end of chromosome arm 2AS (Fig. 2a). This introgression was previously described as “2NS” and originates from *Ae. Ventricosa* (Walkowiak et al. 2020). It contains a source of wheat blast resistance, as well as a gene cluster containing the rust resistance genes *Lr37*, *Yr17* and *Sr38* (Bariana and McIntosh 1993; Helguera et al. 2003; Cruz et al. 2016). To verify that the introgression detected in our analysis corresponds to the 2NS translocation, we mapped whole-genome sequencing (WGS; Walkowiak et al. 2020) reads onto the genome of cultivar Renan (Aury et al. 2022), which, based on our analysis, contains the 2NS introgression. Indeed, the *Ae. ventricosa* reads show a coverage maximum at the position of the suspected introgression, confirming previous reports of cv. Renan carrying this introgression (Supp. Fig. 1b, Aury et al. 2022).

An introgression with unclear origin common to 54 accessions is present in chromosome arm 1DL with a size of ∼11.6 Mb. Comparison of 1D chromosomes with reference-quality assemblies (RQAs), showed that wheat cv. Jagger has an abrupt loss of sequence collinearity with CS at around position ∼487 Mb (Fig. 2c). To investigate the origin of the introgression found in the WHEALBI accessions, we used genotyping-by-sequencing (GBS) data of three accessions predicted to contain the introgression (cv. Crvena Zvezda, Spada and Blueboy, Schulthess et al. 2022) and mapped them to both Jagger and CS. The GBS data confirmed that Crvena Zvezda, Spada and Blueboy have an introgression at the distal end of chromosome 1DL starting at ∼487 Mb (Fig. 2c, Supp. Fig. 2) when using CS as reference. However, when mapping the GBS data to Jagger, we found no reduced sequence coverage in this region, indicating that all four cultivars share the same introgression. Additionally, Illumina reads from *T. timopheevii* (Walkowiak et al. 2020) mapped at a high coverage in the region of the candidate introgression in Jagger (Fig. 2c), which is in line with the findings of a recent preprint (https://doi.org/10.1101/2023.10.04.560829). From this, we conclude that the donor of the 1D introgression is *T. timopheevii* or a species closely related to it.

A previously described introgression with an unknown donor (Cheng et al. 2019; Kale et al. 2022; Schulthess et al. 2022) is present in 101 accessions (54 of which are current varieties). This introgression of ∼30 Mb is located in chromosome 5B. Interestingly, the size of the introgressed fragment varies between accessions. The longest version was found in cv. Renan, which is 30 Mb (Fig. 2d, replacing a 37.5 Mb segment in CS). This introgression is also present in the chromosome-scale assemblies of Fielder, Mace, Jagger, Stanley and Attraktion. Notably, they have two different versions of the introgression, both shorter than the version found in Renan (Fig. 2d, Supp. Fig. 3).

To identify potential donor species for the introgression, we mapped Illumina reads from *T. timopheevii*, *Ae. ventricosa*, *T. ponticum* and *Ae. speltoides* against the genomes of Renan and Attraktion. WGS reads originating from *T. timopheevii* showed a high sequence coverage in the putative introgression regions for Renan and Attraktion (Fig. 2d) which was not the case for the other wheat relatives (Supp. Fig. 4, Supp. Fig. 5). Coincidentally, Attraktion has another introgression which is reported to originate from *T. timopheevii* on chromosome 2B (Walkowiak et al. 2020; Kale et al. 2022). We made use of this and compared the coverage of *T. timopheevii* reads between the introgression of chromosome 2B and chromosome 5B found in Attraktion (Supp. Fig. 6). The median *T. timopheevii* read density in the introgressions on chromosome 2B and chromosome 5B is highly similar (61,723 vs. 61,705 reads/Mb), and much higher than in the rest of the genome (median = 4,246 reads/Mb). This strongly suggests that the introgression on chromosome 5B comes from a closely related (if not the same) donor species as the one on chromosome 2B, which is accepted to stem from *T. timopheevi*. The 5B introgression in Renan contains 307 genes including two NB-ARC LRR (NLR) resistance gene analogs. One of them lies in a cluster of 24 Auxin responsive SAUR genes, which were previously implied to show a role in drought tolerance (He et al. 2021). A full list of genes found in this introgression can be found in Supp. Table S3.

While the more frequent introgressions described above are often documented sources for one or multiple known beneficial traits in the wheat gene pool, this is not known for rare introgressions. We found such introgressions for example in the accession Meteor on chromosome 5D spanning ∼21.5 - 33 Mb in CS, on 6D in the accession 122348 (∼483 - 495.3 Mb) and on 7D in the accession Precoce (telomere - 11.5 Mb, Fig. 2e). In Chinese Spring the segments replaced by these rare introgressions contain 9 (Meteor_5D), 50 (122348_6D) and 81 (Precoce_7D) NLR genes. As introgressions often replace homoeologous segments, it is conceivable that these introgressions introduce new NLR diversity, originating from wild relatives into the wheat gene pool.

### An introgression in cv. Pamukale contains a QTL for powdery mildew resistance

Another example of a rare, previously undescribed introgression is found on chromosome 7D, private to cv. Pamukale (WW-444). Since previously described introgressions were often linked with pathogen resistance, for example the 1RS.1BL translocation (Crespo-Herrera et al. 2017), we considered Pamukale as an interesting cultivar for mapping novel powdery mildew resistance (*R*) genes. Phenotyping on Pamukale with 66 wheat powdery mildew (*Bgt)* isolates from a worldwide collection (Sotiropoulos et al. 2022) identified resistance to 24 *Bgt* isolates from different regions of the world (Fig. 3a). To map the underlying *R* locus, we crossed Pamukale with cv. Frisal, which is broadly susceptible to *Bgt* at the seedling stage (Fig. 3b). An F2 population inoculated with isolate Bgt_10001 segregated 34 resistant: 99 susceptible (chi^2^_1:3_ = 0.02; P_1df_ = 0.88) indicating that resistance was conferred by a single recessive locus. In a separate test with *Bgt* isolate Bgt_07004, similar results were obtained, with 40 F2 individuals which were scored resistant (Supp. Table S4). The difference in ratio was attributed to error or quantitative variation in phenotyping.

**Fig. 3.**
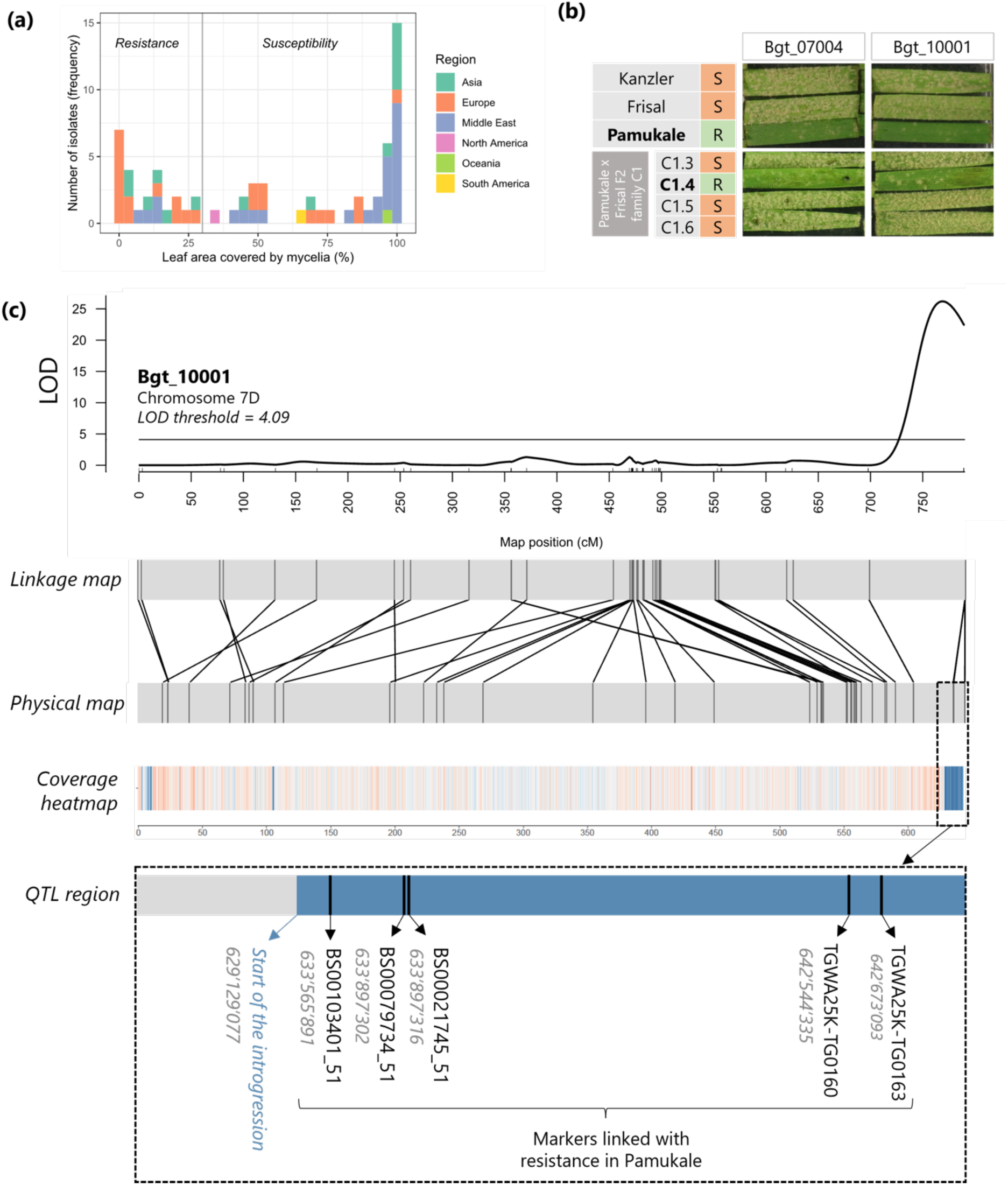
A QTL for powdery mildew resistance in cv. Pamukale is located within an introgression in chromosome 7D. **(a)** Frequency of resistant and susceptible phenotype against 66 *Bgt* isolates from a worldwide collection. A leaf coverage below 30% corresponds to a resistant phenotype, while above 30% is considered susceptible. **(b)** Pamukale provides seedling stage resistance against the *Bgt* isolates Bgt_07004 and Bgt_10001. The F2 progeny individuals (lower panel) show a recessive segregation of the resistance phenotype. The phenotype was evaluated eight days after spray inoculation of detached leaf segments. **(c)** A QTL locus for powdery mildew resistance in chromosome 7D is linked with five SNP markers located in an alien introgression unique to the cultivar Pamukale. The representation of the *R* gene locus is not in scale.

All Pamukale x Frisal F2 individuals were genotyped with a 25K SNP wheat array and a linkage map with 2,058 polymorphic markers distributed across the whole genome was constructed (Supp. Table S5). The linkage map was used for QTL mapping, which revealed a QTL for powdery mildew resistance to both *Bgt* isolates, located at the end of the long arm of chromosome 7D (Fig. 3c, Supp. Fig. 7), colocalizing with the 7D introgression unique to Pamukale. The underlying gene in the QTL was named *PmPam*.

The QTL was flanked by the markers GENE-4442_121 and BS00103401_51, with the latter being completely linked with *PmPam* (LOD > 20; Fig. 3c). There were four additional SNP markers located in the same position as BS00103401_51 in the linkage map, which were perfectly linked (except for a few missing SNP marker calls). Interestingly, we observed that they were physically apart from each other, ranging from 633,565,891 to 642,673,093 in the CS reference genome (Fig. 3c). This clustering in the linkage map was attributed to the presence of the introgression in Pamukale (starting at position ∼629.1 Mb) suppressing recombination in that genomic region. We therefore assume the QTL spanned the entire 14 Mb of the introgression, making fine mapping of *PmPam* nearly impossible (Fig. 3c).

*Pm5* is a recessive *R* allelic series located at the end of chromosome 7B (Xie et al. 2020) and has homoeologs in chromosomes 7A and 7D. To exclude that *PmPam* was derived from the *Pm5* locus, we performed a synteny comparison between the ends of chromosomes 7A, 7B and 7D, and found that in chromosome 7D the *Pm5* homoeolog is located at 616,968,098 in the CS genome (Supp. Fig. 8). It is therefore not in the introgression region, which starts at 629,129,077, and we concluded that *PmPam* is distinct from *Pm5* and its homoeologs.

### Multiple wheat accessions show evidence for inter-homoeolog recombination

We found chromosomal segments in multiple accessions with a relatively even coverage ratio of ∼2, but there was a concomitant absence of reads in the corresponding segment on one of the homoeologous chromosomes. One example is the modern variety Ritmo (WW-150, Fig. 4a) where the first ∼39 Mb of chromosome 1A have almost no read coverage (median=0.03) whereas the first ∼41 Mb of chromosome 1D have a median coverage ratio of 1.87. This indicates that the first ∼39 Mb of chromosome 1A in Ritmo are missing, while the beginning of chromosome 1D is duplicated. An explanation of such an observation is a reciprocal translocation between homoeologous chromosomes. Reciprocal translocations were also described in rice (Wicker et al. 2015), synthetic allopolyploid wheat (Zhang et al. 2020) and more recently in newly produced *Am. muticum*/hexaploid wheat introgression lines (Coombes et al. 2023). In the case of Ritmo we propose that chromosome 1D is ‘regular’, whereas the distal end of chromosome arm 1AS was replaced by the first 41 Mb of chromosome 1D, which would therefore occur twice in the genome of Ritmo (Fig. 4)

**Fig. 4.**
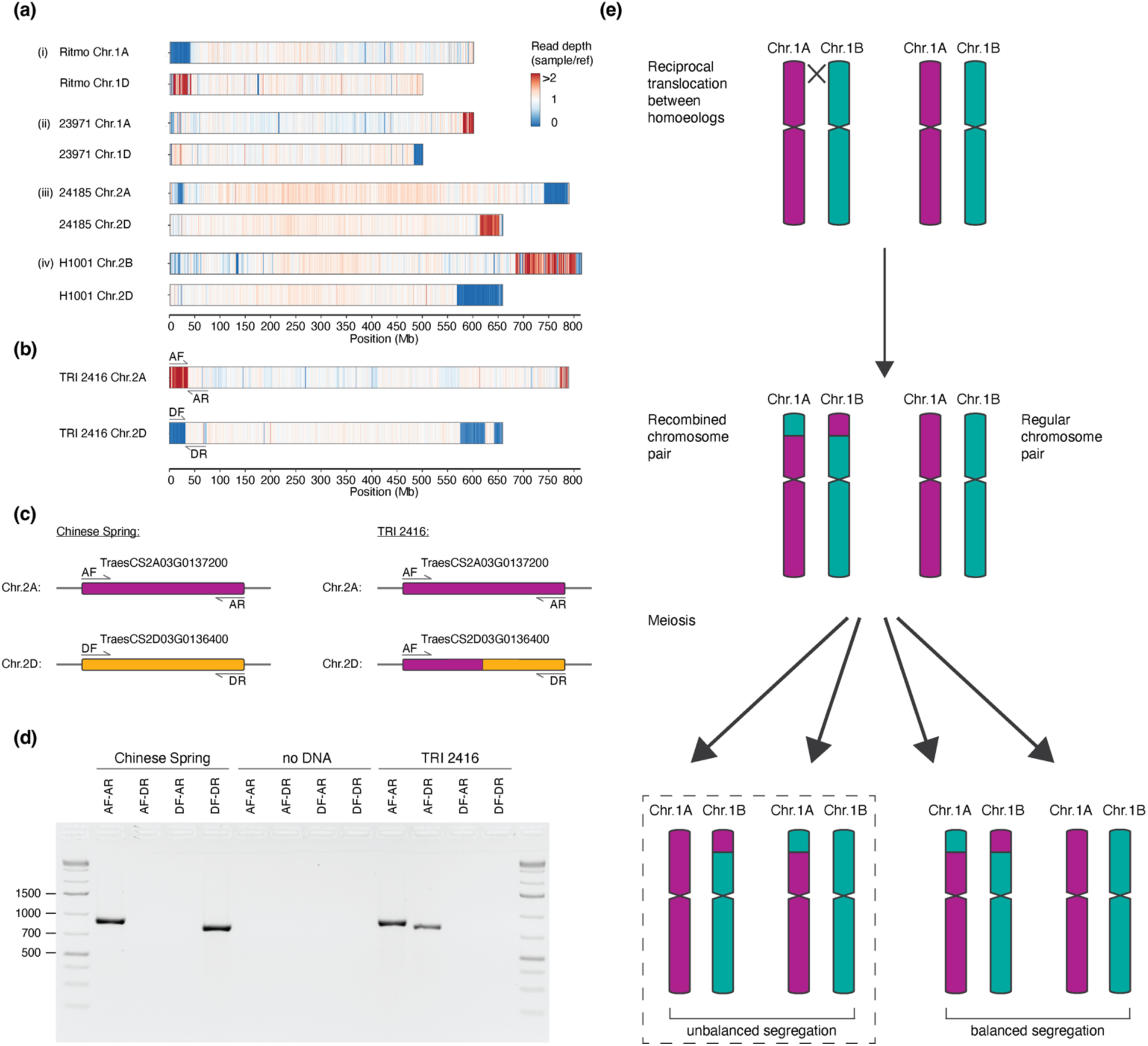
Evidence for recombination between homoeologous chromosomes in some genotypes of bread wheat. **(a)** and **(b)** Examples of putative inter-homoeolog recombination events. The heatmap shows coverage of normalised exome capture reads mapped against CS. Values are calculated in 500 kb bins. An example used for PCR verification. Location of primers are indicated and shown in more detail in **(c)** Scheme of primers designed to confirm inter-homoeolog translocation/duplication by PCR across the translocation breakpoint. The break point lies within the homoeologous genes *TraesCS2A03G0137200* and *TraesCS2D03G0136400*. (**d**) 1.2% Agarose gel loaded PCR products. There is a PCR product of the expected size for the primer combinations AF-AR and DF-DR in CS. The DF-DR product is absent in TRI 2416, and there is amplification for AF-DR **(e)** Schematic model of reciprocal translocations between homoeologs. Translocation is shown with the example of chromosomes 1A and 1B. After recombination, meiosis leads to the segregation of homoeologous chromosomes, in which two gametes contain unbalanced chromosomal content.

Another example was found in the Albanian traditional cultivar TRI 2416 (WW-078, Fig. 4b), where the first 36 Mb of chromosome 2A have a median coverage ratio of 2.208, whereas the first 31 Mb of chromosome 2D have almost no mapped reads (median = 0.017). We propose that chromosome 2A is ‘regular’, while the distal end of chromosome arm 2DL was replaced by the first 36 Mb of chromosome 2A, which therefore occurs twice in the genome of TRI 2416. To confirm this rearrangement, we first identified the putative translocation breakpoints bioinformatically by identifying regions where Illumina reads map on chromosome 2A, and their mate reads map on chromosome 2D. We identified the candidate break points to be in the homoeologous genes *TraesCS2A03G0137200* (chromosome 2A) and *TraesCS2D03G0136400* (chromosome 2D respectively) (Supp. Fig. 9). Indeed, we found sequencing reads covering the recombination breakpoint (Supp. Fig. 9). We then designed primers, based on the CS genome, and performed PCR amplifications across the breakpoint, that confirmed the proposed arrangement (Fig. 5b). This indicates that the breakpoint of the 2A-2D translocation found in TRI 2416 indeed lies in the genes *TraesCS2A03G0137200* and *TraesCS2D03G0136400*. Based on 2A- and 2D-specific SNPs we postulate the breakpoint to lie between the genomic positions 35,685,613 and 35,685,702 bp of chromosome 2A (31,143,922 - 31,144,011 bp in chromosome 2D, Supp. Fig. 9b).

**Fig. 5.**
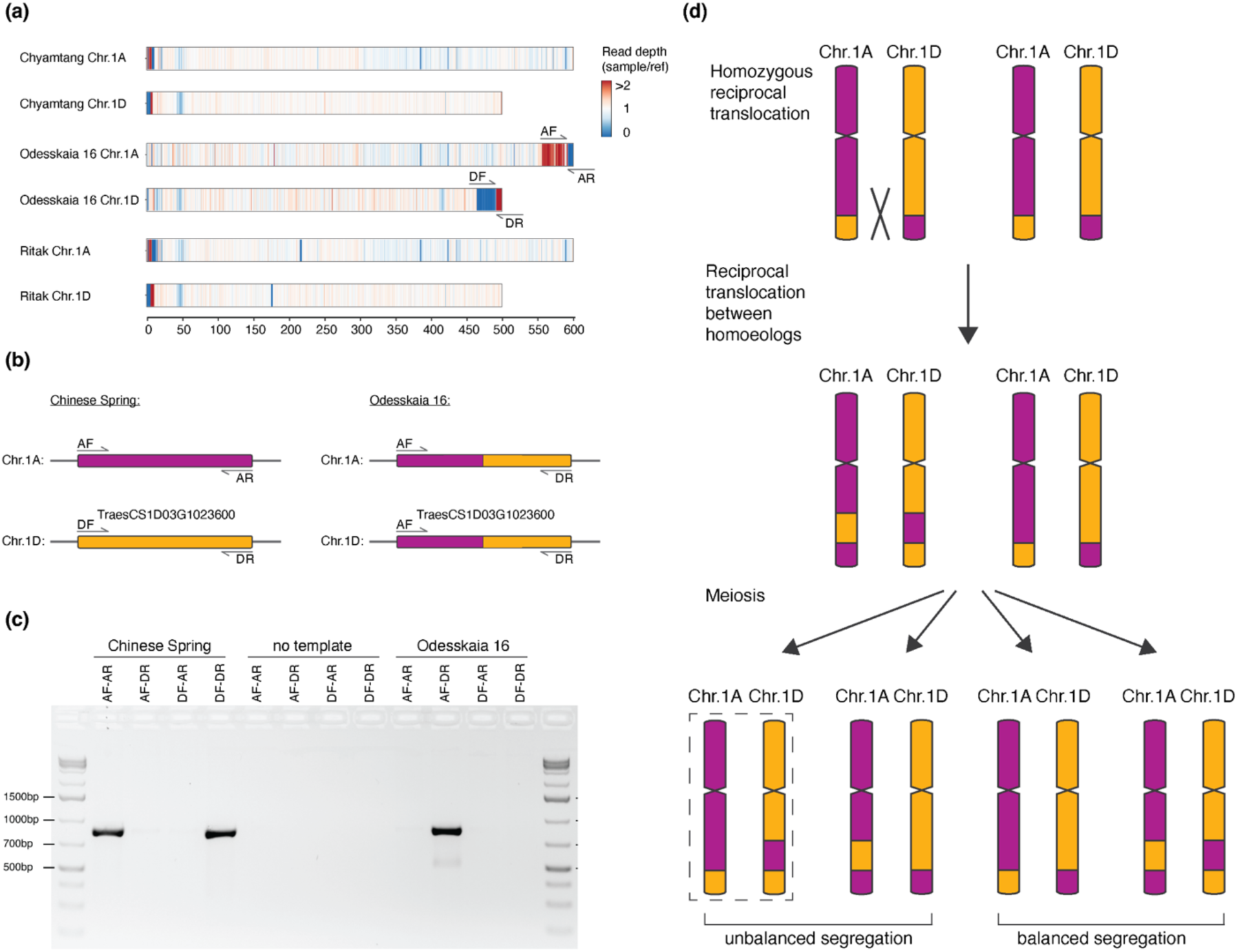
Evidence for recurrent recombination between homoeologous chromosomes in some genotypes of bread wheat **(a)** Examples of suspected double inter-homoeolog recombination events. Heatmap showing coverage of normalised exome capture reads mapped on the CS genome. Values are calculated in 500 kb bins. Location of primers are indicated and shown in more detail in: **(b)** Scheme of primers designed for PCR amplification across the suspected translocation break point, which was within gene *TraesCS1D03G1023600*. **(c)** 1.2% agarose gel loaded with PCR products. **(d)** Schematic model of reciprocal translocations between homoeologs, starting with a heterozygous, single inter-homoeolog translocation carrier. Translocation is illustrated between chromosomes of the A and D genomes. Further recombination generates recombined chromosomes. Chromosome structure, as seen in Odesskaia 16 is shown with a dotted rectangle.

Based on these results, we propose a model for inter-homoeologous exchange similar to what has been proposed in rice (Wicker et al. 2015, Fig. 4e): the first step is a symmetric reciprocal translocation between homoeologous chromosomes (1A and 1B in the example in Fig. 4e). At meiosis, there are then four possible gametes, two with balanced chromosome content, and two with a duplication of one homoeolog sequence and a loss of the other. Descendants of these cell lines can then, in later generations, become homozygous.

Interestingly, we also found cases where a subtelomeric region shows coverage ratio ∼2, while the adjacent telomeric segment has a coverage ratio of ∼0, with the pattern being inverted on one of the homoeologous chromosomes (Fig. 5a). This is especially prominent in the accession Odesskaia 16 (WW-403), where a 38 Mb (554 - 592 Mb) segment on the long arm of chromosome 1A shows a median coverage ratio of 2.08, and the adjacent (telomeric) 6.5 Mb have a median coverage ratio of 0.015. A homoeologous segment on chromosome 1D (464 - 491 Mb) shows a coverage ratio of 0.027, whereas the telomeric 7.5 Mb has a coverage ratio of 2.37 (Fig. 5a). Expanding the model of inter-homoeologous exchange, we propose that these examples resulted from two recombination events (Fig. 5b): the evolution of the Odesskaia 16 lineage started having a homozygous translocation in which the telomeric 7.5 Mb of chromosome 1D and the telomeric 38 Mb of chromosome 1A were swapped (Fig. 5d). Subsequently, a second, larger inter-homoeolog recombination exchanged the telomeric ∼50 Mb between the two recombinant chromosomes. As for the model in Fig. 5d, unbalanced segregation led to the duplication/deletion pattern observed in Odesskaia 16.

We again used information from paired-end Illumina reads to identify the precise translocation breakpoints. Although the proximal breakpoint could not be identified, either because the gene containing the breakpoint was not sequenced using exome capture (due to only 59.8% of genes being sequenced or because the breakpoint does not lie within a gene), we could estimate, based on sequence coverage data, the breakpoint to lie between the region corresponding to 463,719,382 and 463,727,141 in CS chromosome 1D. The distal breakpoint was found in the gene *TaesCS1D03G1023600*. Again, we found forward and reverse read pairs covering the recombination breakpoint (Fig.6). This chromosomal position had double the coverage on chromosome 1A proximal to the breakpoint, and no coverage distal of the breakpoint on and, complementary, no coverage on chromosome 1D before the breakpoint and double the coverage after the breakpoint (Fig. 6a). We used homoeolog-specific PCR to confirm this breakpoint; The primer combinations AF-AR and DF-DR gave a PCR product of the expected length (832 bp), when using genomic DNA of accession CS (Fig. 5c). In the case of Odesskaia 16 there is only a PCR product for the primer combination AF-DR.

**Fig. 6.**
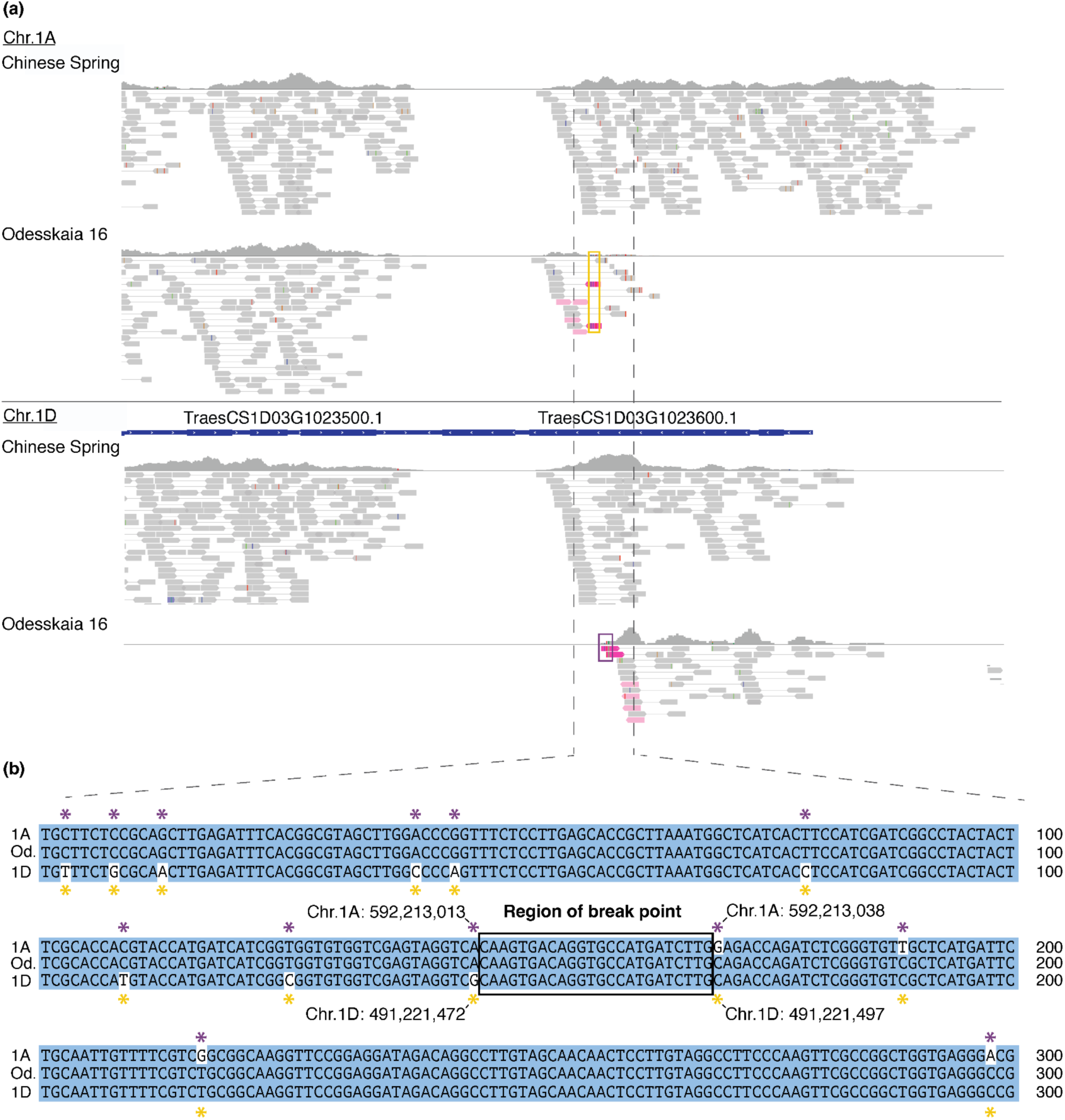
Identification of inter-homoeolog recombination break point. **(a)** Visualization of mapped reads from Chinese Spring and Odesskaia 16 on CS exported from IGV (Robinson et al. 2011). Mate-pairs going across the break point (one mate mapping to Chr.1A the other to Chr.1D) were colored in light pink. Reads that show both 1A and 1D specific SNPs in the same read (i.e. chimeric reads) are colored in pink and framed by either yellow or purple rectangles. **(b)** Alignment of Chr.1A and Chr.1D in the region surrounding the suspected break point 1A or 1D specific SNPs are indicated by asterisks.

From these data, we conclude that the WHEALBI collection contains multiple wheat accessions which contain duplicated chromosome segments that resulted from inter-homoeolog recombination followed by unbalanced segregation (Fig. 5d).

## Discussion

In this study, we exploited the public resource of exome capture data from a worldwide collection of 434 hexaploid wheat accessions, which includes currently used varieties, old cultivars, and landraces. Although the exome capture data only represented about 59.8% of all genes and less than 1% of the genome, the data proved to be highly useful for the automated identification of introgressed chromosomal segments. Using the straightforward approach of analysing sequence read coverage normalised by reads of a reference, we shed light on the diverse introgression landscape found in the WHEALBI collection (Pont et al. 2019), composed of over 26,664 introgressions.

Wheat breeding depends on diversity in the gene pool. The very high number of candidate introgressions identified here indicates that a considerable part of the diversity in the wheat gene pool derives from such events. While some of the introgressions were made by humans, many may in fact be the result of substantial and natural gene flow.

These numbers are congruent with findings of a previous study that identified hundreds of introgressions based on 10 RQAs (Walkowiak et al. 2020), but in this study we went a step further, providing an introgression atlas of wheat material used in current breeding programs. Nonetheless, the numbers presented in this study are likely an overestimation, because in our method regions of low coverage must be continuous. This means that for example, if an internal segment of an introgression recombines and is replaced by common wheat, the internal segment will have a normal coverage ratio of ∼1 and the introgression will therefore be split in two (Zhou et al. 2020). Our method will then count this as two introgressions, although it came from a single event.

The chosen approach allowed us to assess the frequencies of previously known major introgressions in the wheat gene pool. We could, for example, confirm the prevalence of an introgression from *Ae. markgrafii* found on chromosome 2D in western breeding material (Walkowiak et al. 2020; Keilwagen et al. 2022). Strikingly, more than half of all accessions had multiple introgressed segments of 10 Mb or larger, with smaller introgressions being even more prevalent. The data from introgressions that are shared between multiple accessions indicates that introgressed segments are often partially removed, likely through back-crosses with lines which do not contain them (Zhou et al. 2020). One example is the 5B introgression shown in Fig. 2d. Therefore, it is often not possible to determine whether an introgression happened only once and was later partially lost, or whether a similar fragment was introgressed multiple times. Despite the difficulties in estimating how frequently alien chromosome segments are introgressed, our data show that almost all wheat accessions studied here contain a complex mosaic of chromosomal segments coming from different genetic backgrounds.

### Diverse donor species contribute potentially important genetic material

We could identify the potential donor species for a few major introgressions. This is, in principle, only possible if genomic sequence data of the donor species is available. Indeed, we confirmed *Ae. ventricosa* as the donor species for the previously described introgression on chromosome 2A (Walkowiak et al. 2020; Keilwagen et al. 2022). More importantly, we identified *T. timopheevii* as the putative donor of a previously described 5B introgression (Cheng et al. 2019; Kale et al. 2022; Schulthess et al. 2022). We used publicly available Illumina reads from *T. timopheevii*, *Ae. ventricosa*, *Th. ponticum* and *Ae. speltoides*, which showed that *T. timopheevii* is the best candidate donor because its reads mapped much more frequently and with higher quality to wheat accessions which contain the introgression. However, since read mappings were not perfect, it is possible that the introgression came from a different haplotype or a close relative of *T. timopheevii*. This emphasises the need for additional genomic sequencing of wild wheat relatives, so that donors of important genes can be identified more readily. However, even without donor sequence data, it is feasible to infer to some degree potential donor species simply based on the level of sequence homology.

A particularly important finding of this study was that numerous introgressions were found in single wheat accessions. Indeed, we identified 118 introgressions of 10 Mb or larger unique to single accessions in the collection. We are aware that the WHEALBI collection covers only a small fraction of the many thousands of wheat accessions that comprise the worldwide gene pool. Nevertheless, our data indicate that the wheat gene pool contains high numbers of rare but potentially valuable introgressions which must be first identified and then linked to agronomically important genes. We demonstrated the value of a specific introgression through QTL mapping, where we identified *PmPam*, a powdery mildew resistance gene originating from an introgression on the long arm of chromosome 7D found exclusively in the Turkish cv. Pamukale. This major QTL segregates recessively and is the second recessive *Pm*-QTL reported on chromosome 7D, the other being *WTK4,* which is located on the short arm of chromosome 7D (Gaurav et al. 2021; Tang et al. 2023). The QTL identified in Pamukale confers resistance to multiple *Bgt* isolates originating from different geographical regions. This makes it a potentially valuable resource in resistance breeding. Moreover, the introgression containing *PmPam* is probably rare, as it is present in a single cultivar of the WHEALBI collection, and this might represent an additional advantage. In fact, broadly used, introgressed resistance genes are often overcome due to extended and strong selection pressure against avirulent pathogens (Kunz et al. 2023), which further emphasise the need of mapping new and rare resistance genes. Our finding illustrates that rare introgressions already present in the breeding pool, can harbor unexploited, beneficial traits that can contribute to sustainable agriculture. This highlights the usefulness of introgression catalogues such as the one presented here.

### Inter-homoeolog recombination is frequent in the wheat gene pool

Our analysis of sequence coverage also revealed compelling evidence for recombination between homoeologous chromosomes in at least 55 accessions (Supp. Table S6). Of these, at least eight do not seem to be independent, as they share one inter-homoeolog-recombination breakpoint, and all of them are either old cultivars or cultivars from Nepal (Supp. Fig. 10). Hexaploid wheat behaves as a genetic diploid since chromosomes of different subgenomes are assumed not to recombine. This bivalent behaviour is enforced by several genes among which the *Ph1* locus is the most important (Rey et al. 2017). It was suggested that inter-homoeologous recombination depends on the suppression of *Ph1* (Coombes et al. 2023). Our data suggest that despite this tight genetic control, inter-homoeolog recombination can occur in hexaploid wheat. While we identified most cases in landraces (14) and old cultivars (20), we also found evidence for such an event in the modern varieties such as Ritmo (Fig. 4a), Dekan, KWS Magic and Palesio (Supp. Fig. 11). Suppression of *Ph1* is used to enable chromosome pairing in introgression breeding (Martín et al. 2017). Therefore, some of the observed inter-homoeologous recombination events may be a by-product of introgression breeding. However, most of the events reported here were in old cultivars and landraces, which indicates that inter-homoeolog recombination also occurs naturally.

Previous studies showed that recombination between homoeologous chromosomes can occur. One study documented inter-homoeolog recombination in artificial tetraploid wheat (Zhang et al. 2020) and found recombination to occur primarily in low-copy sequences (i.e. genes). In cases where we could precisely identify the recombination breakpoints, we also found them inside orthologous genes, indicating that inter-homoeolog chromosome pairing occurs in syntenic positions. Similar findings were also made in rape seed and wheat using transcriptome data (He et al. 2017).

It must be emphasised that we identified these cases, because the inter-homoeolog recombination event was followed by unbalanced segregation, that resulted in recombinant segments being duplicated in a given genotype. A simple (symmetrical) exchange of chromosomal segments between homoeologs would not have been detectable purely based on sequence read coverage. It is therefore possible that the WHEALBI data set contains such symmetric exchanges between homoeologs which we were not able to detect.

In this context it is interesting that we also found cases that presumably resulted from multiple, sequential inter-homoeolog recombination events (Fig. 5). Such events depend on an initial, symmetric, inter-homoeolog recombinant in a homozygous genotype. Further recombination and duplication events led to the observed types (Fig. 5). This would mean that symmetric recombinants must be able to recombine with non-recombinant wheat lines, or they would likely be selected against (either naturally or by humans) and rapidly decline in frequency. Indeed, maintenance of heavily re-arranged chromosomes in the wheat gene pool has been reported previously: for example, the wheat variety Arina, like other European winter wheat lines (Walkowiak et al. 2020), has a major rearrangement where the short arms of chromosomes 5B and 7B are exchanged, but the recombinant Arina can be crossed without restrictions with non-recombinant wheat lines (Walkowiak et al. 2020; Kolodziej et al. 2021). The underlying genetic mechanisms are well known: recombinant chromosomes would recombine with non-recombinant ones by forming cross tetrads, a constellation that allows homologous chromosome segments to pair correctly (Copenhaver et al. 2000; Golczyk et al. 2008). A schematic description of the proposed mechanism is shown in Fig. 7.

**Fig. 7.**
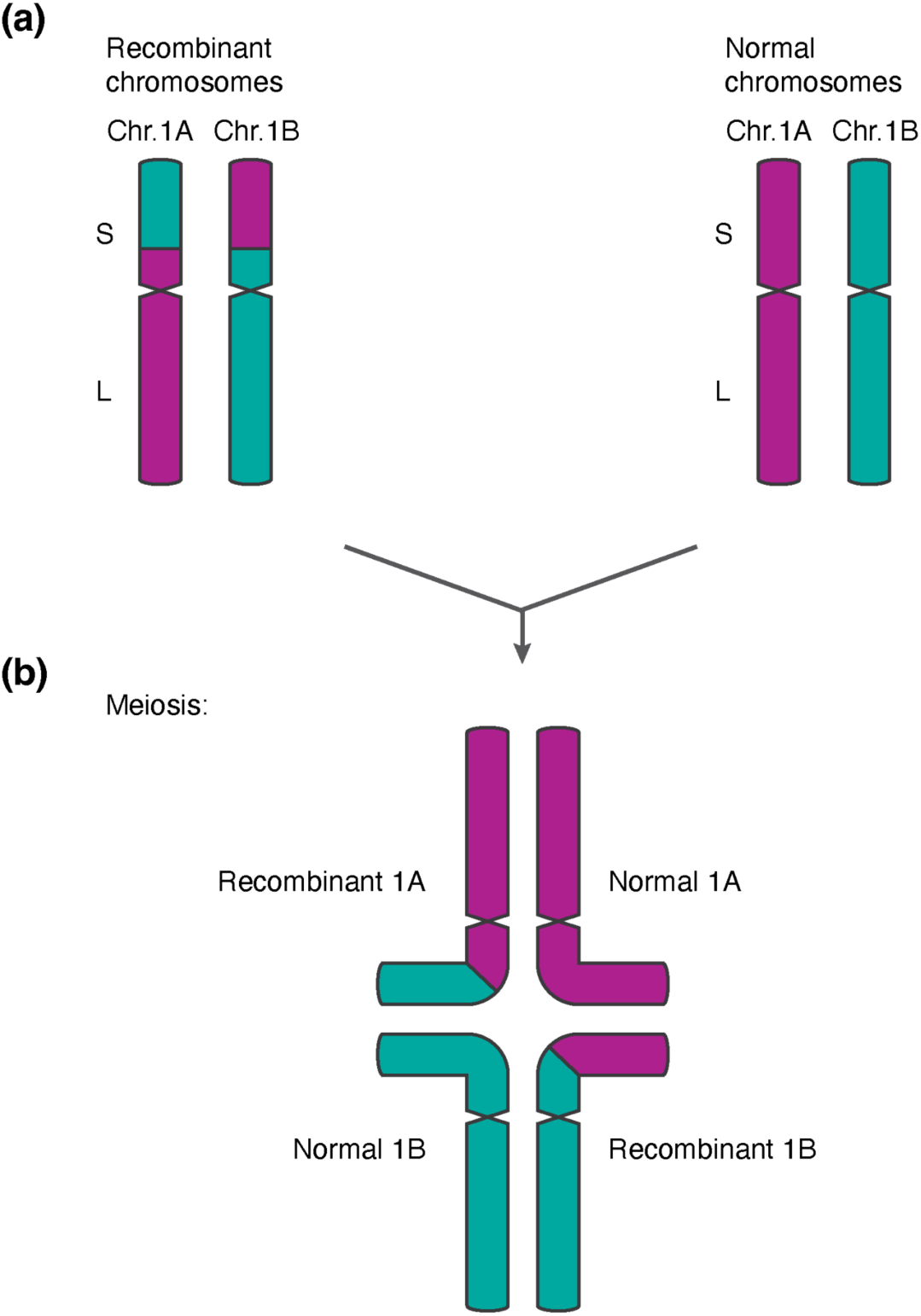
Schematic representation of cross-tetrad formation between inter-homoeolog recombinants and non-recombinants. **(a)** Schematic germline karyotype with one set of homoeologs carrying a balanced reciprocal translocation between homoeologs, and the other being normal. **(b)** Model for pairing of chromatids as cross-tetrads. The recombinant part of the short arm of Chr.1A pairs with the homologous segment in the normal Chr.1B. The non-recombinant parts of Chr.1A pair normally. Commensurately, the recombinant part of the short arm of Chr.1B pairs with the homologous segment in the normal Chr.1A, while the non-recombinant parts of Chr.1B pair normally.

In conclusion, our study confirms that the wheat gene pool comprises a plethora of accessions with complex patterns of chromosomal introgressions and chromosomal rearrangements, and further provides an introgression landscape easily accessible at https://github.com/matthias-heuberger/whealbi_introgression_viewer, providing novel tools for future breeding improvement.

## Acknowledgements

We thank Robert McIntosh for critical reading of the manuscript.

## Author contributions

MH, ZB and TW wrote the initial manuscript with input from BK and JSM. The study was conceived by TW, JSM and BK. MH conducted introgression and inter-homoeolog analyses. EJ and VW performed wheat crossing and contributed to plant maintenance and seed propagation. ZB performed phenotyping, DNA extraction for genotyping, and conducted QTL mapping analysis. TW, GH and MH designed PCR experiments. MH conducted PCR experiments. All authors have read and approved the final manuscript.

## Data availability

Linkage map, genotype and phenotype information of the Pamukale x Frisal F2 progeny are available in Supp. Tables S5 and S4 respectively. Normalized coverage data and introgression assignment are available at: https://zenodo.org/records/10406469. Genome assemblies are publicly available and available for download in the following locations. IWGSC RefSeq v2.1: https://urgi.versailles.inra.fr/download/iwgsc/IWGSC_RefSeq_Assemblies/v2.1/; ArinaLrFor, Claire, Jagger, Julius, Kariega, Lancer, Landmark, Mace, Renan, Stanley, SYMattis: https://plants.ensembl.org/info/data/ftp/index.html; Attraktion: https://www.ebi.ac.uk/ena/browser/view/GCA_918797515.1. The WHEALBI exome capture data is publicly available and can be obtained in NCBI with the BioProject number PRJNA524104. The genomic sequence reads of wheat relatives are publicly available in the following locations: *T. timopheevi*, *A. ventricosa*, *T. ponticum* - NCBI Bioproject number PRJNA544491, *A. Markgrafii* and *A. speltoides* - ENA ID PRJEB49121, *T. monococcum -* ENA ID PRJEB61155. GBS data is publicly available in ENA under the ID PRJEB41976.

An R shiny application that can be used to visualise the coverage ratio data generated for this manuscript is available at: https://github.com/matthias-heuberger/whealbi_introgression_viewer.

## Supplementary Figures

**Supp. Fig. 1.**
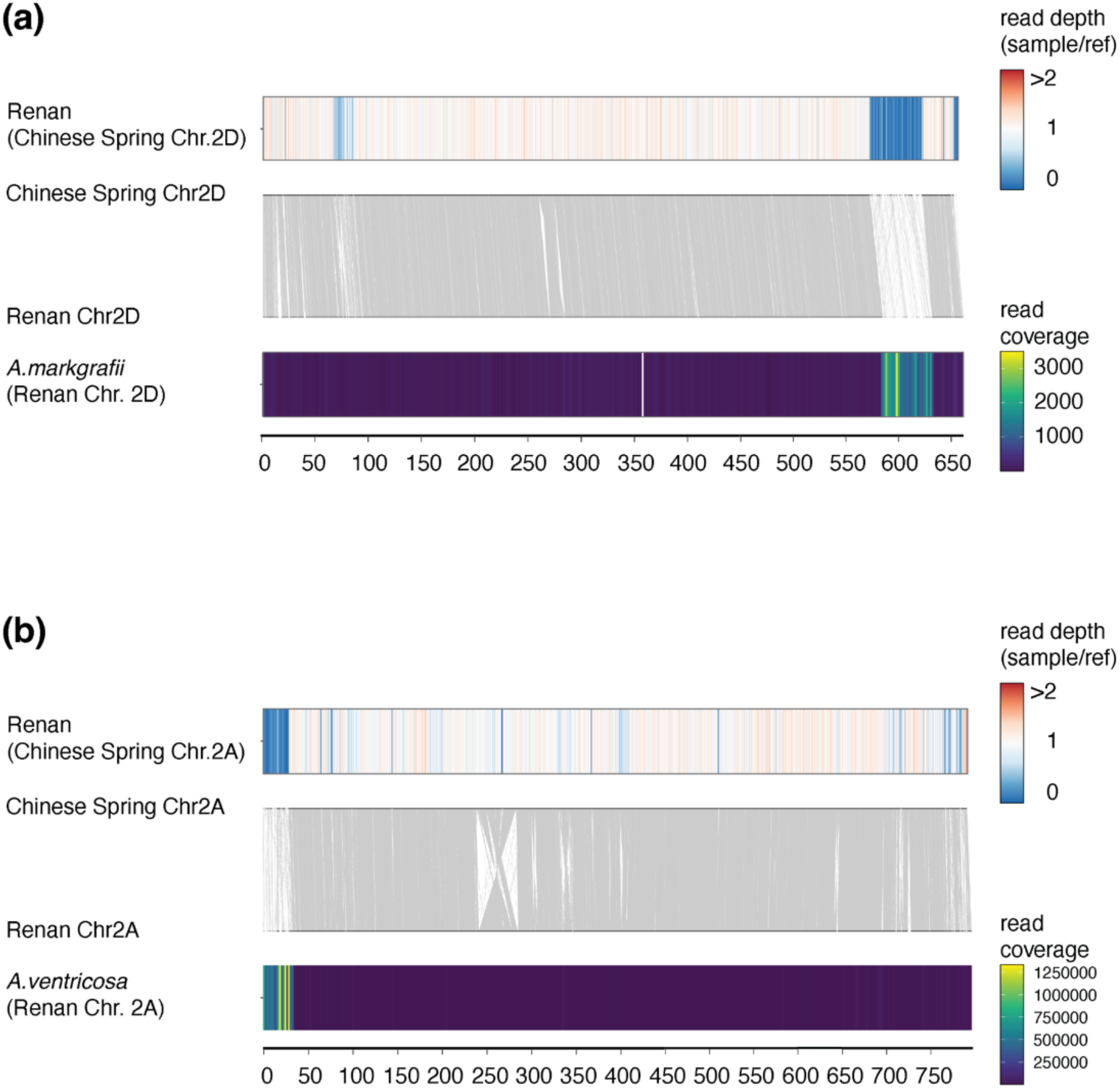
Confirmation of previously described introgressions in Renan **(a)** The top drawing shows the coverage of normalised exome capture reads of Renan on Chr. 2D. The middle drawing shows the chromosome collinearity between Chinese Spring Chr. 2D and Renan Chr. 2D. At the bottom, the read coverage of *A. markgrafii* WGS reads mapped on Renan Chr. 2D is given. Values for exome capture were calculated in 500 kb bins and WGS in 2 Mb bins. **(b)** The top drawing shows the coverage of normalised exome capture reads of Renan on Chr. 2A. In the middle, the chromosome collinearity between Chinese Spring Chr. 2A and Renan Chr. 2A is shown and at the bottom, the read coverage of *A. ventricosa* WGS reads mapped on Renan Chr. 2D given. Values for exome capture were calculated in 500 kb bins and WGS in 2 Mb bins.

**Supp. Fig. 2.**
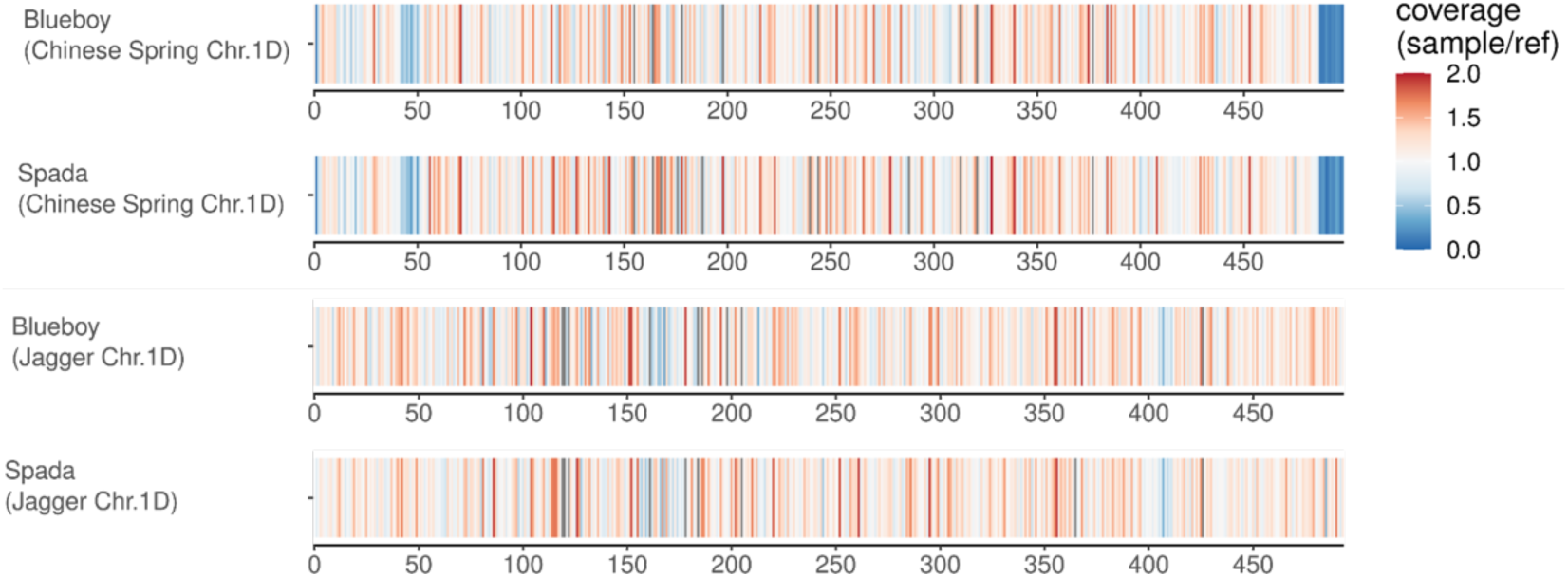
Blueboy and Spada contain the same introgression as Jagger. GBS data of Blueboy and Spada mapped on Chinese Spring Chr. 1D and Jagger Chr. 1D. At the distal end of Chinese Spring Chr. 1D Blueboy and Spada show low coverage, while they show normal coverage at the distal end of Chr. 1D of Jagger. Values were calculated in 2 Mb bins and normalised by GBS read coverage of the respective reference genotype against which the reads were mapped.

**Supp. Fig. 3.**
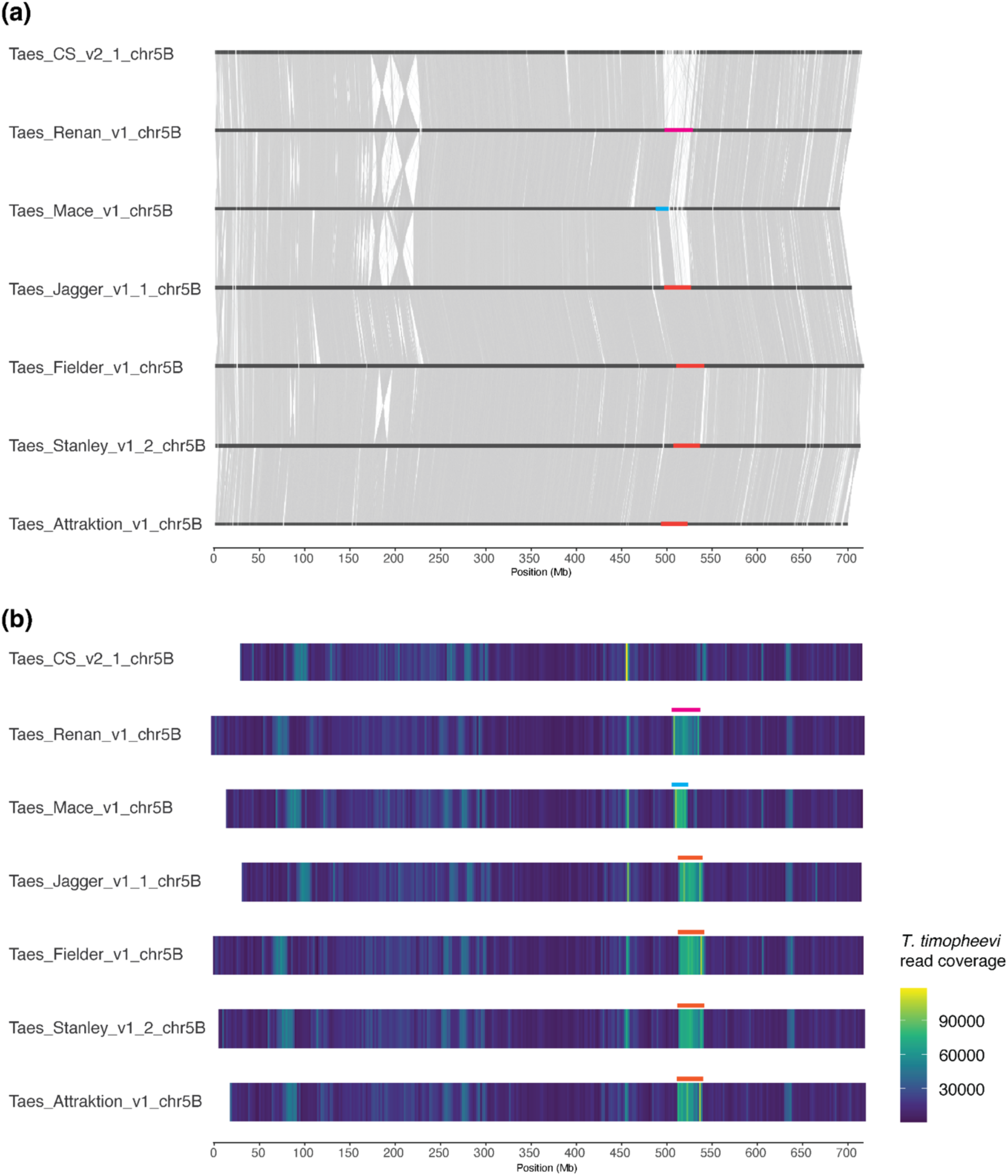
Reference-quality genome assemblies (RQAs) have three different versions of the 5B introgression. **(a)** Chromosome collinearity for Chr. 5B between Chinese Spring and six RQAs containing the 5B introgression. Chromosomes were arranged according to similarity in the region of the introgression. **(b)** *T. timopheevi* WGS heat map showing read coverage of reads mapped on Chr. 5B of Chinese Spring and six RQAs containing the 5B introgression. Values were calculated in 2 Mb bins. To allow visual comparison of the introgression region, the chromosomes were aligned based on collinearity just before the introgression. Scale at the bottom is aligned to Chr. 5B of Fielder.

**Supp. Fig. 4.**
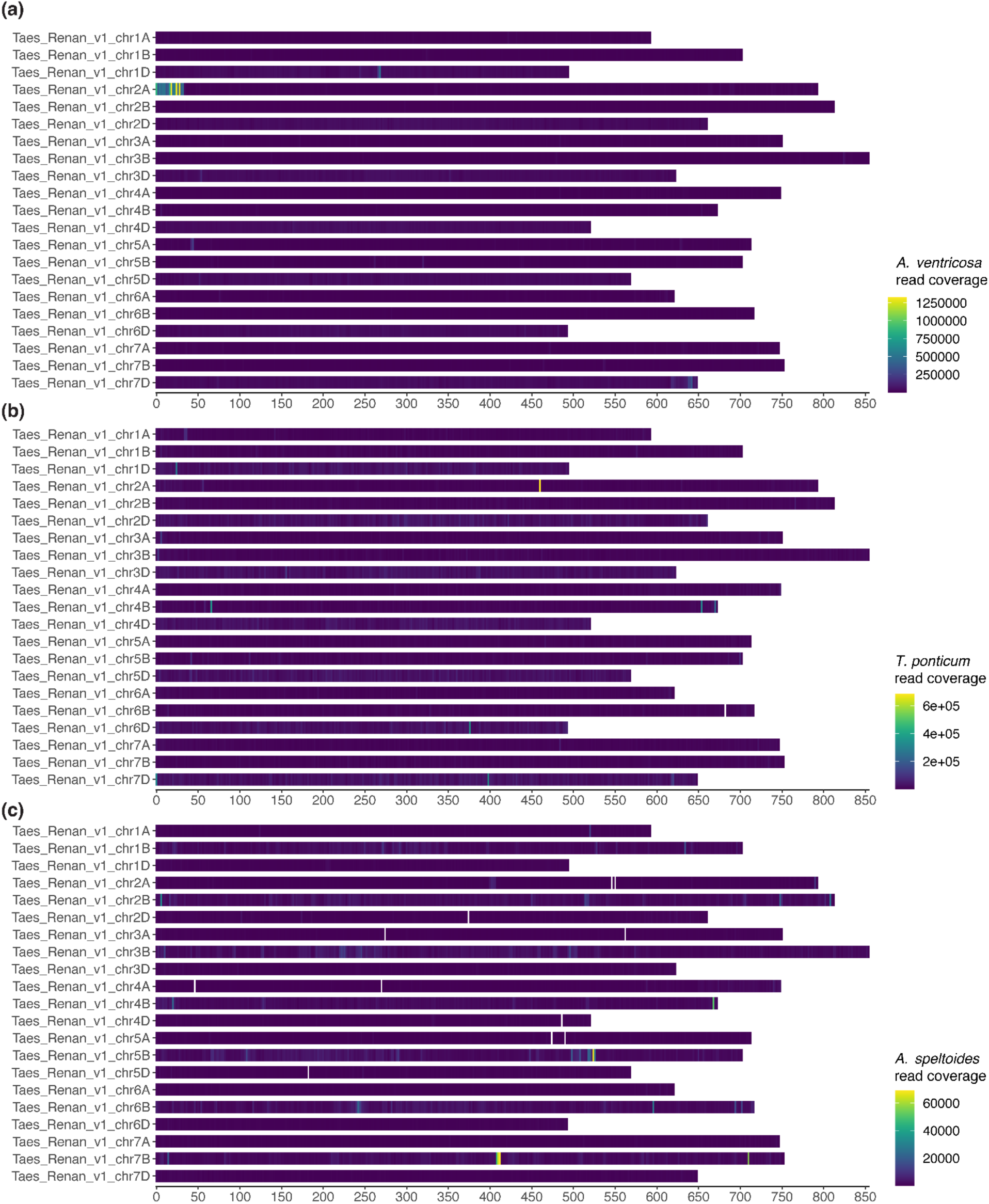
Mapping of wild wheat relatives onto the Renan genome assembly **(a)** Read coverage of WGS reads from *A. ventricosa* mapped on Renan. **(b)** Read coverage of WGS reads from *T. ponticum* mapped on Renan. **(c)** Read coverage of WGS reads from *A. speltoides* mapped on Renan. Values were calculated in 2 Mb bins.

**Supp. Fig. 5.**
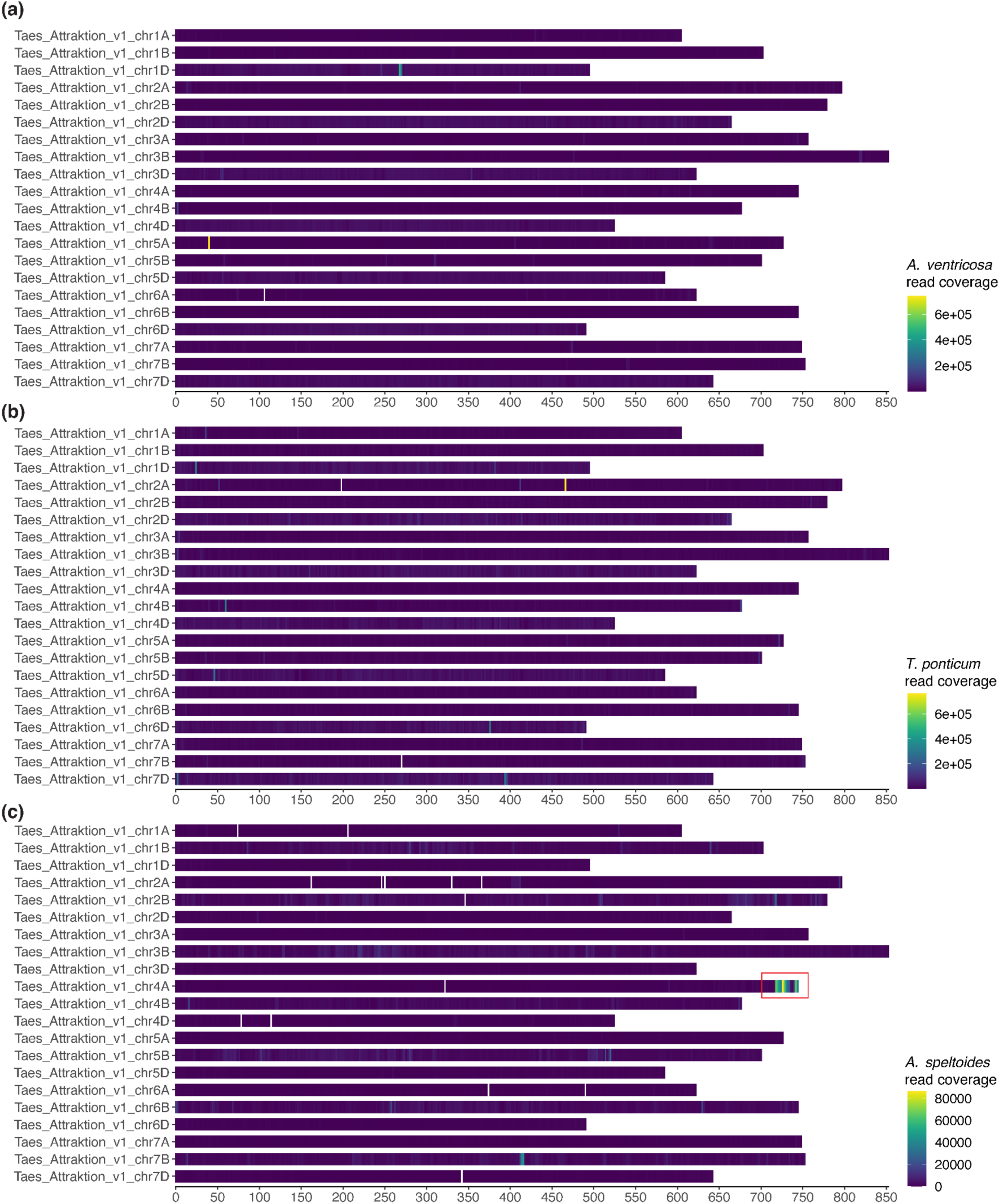
Mapping of wild wheat relatives onto the Attraktion genome assembly **(a)** Read coverage of WGS reads from *A. ventricosa* mapped on Attraktion. **(b)** Read coverage of WGS reads from *T. ponticum* mapped on Attraktion. **(c)** Read coverage of WGS reads from *A. speltoides* mapped on Attraktion. Known *A. speltoides* introgression (Keilwagen et al. 2022) highlighted by red rectangle. Values were calculated in 2 Mb bins.

**Supp. Fig. 6.**
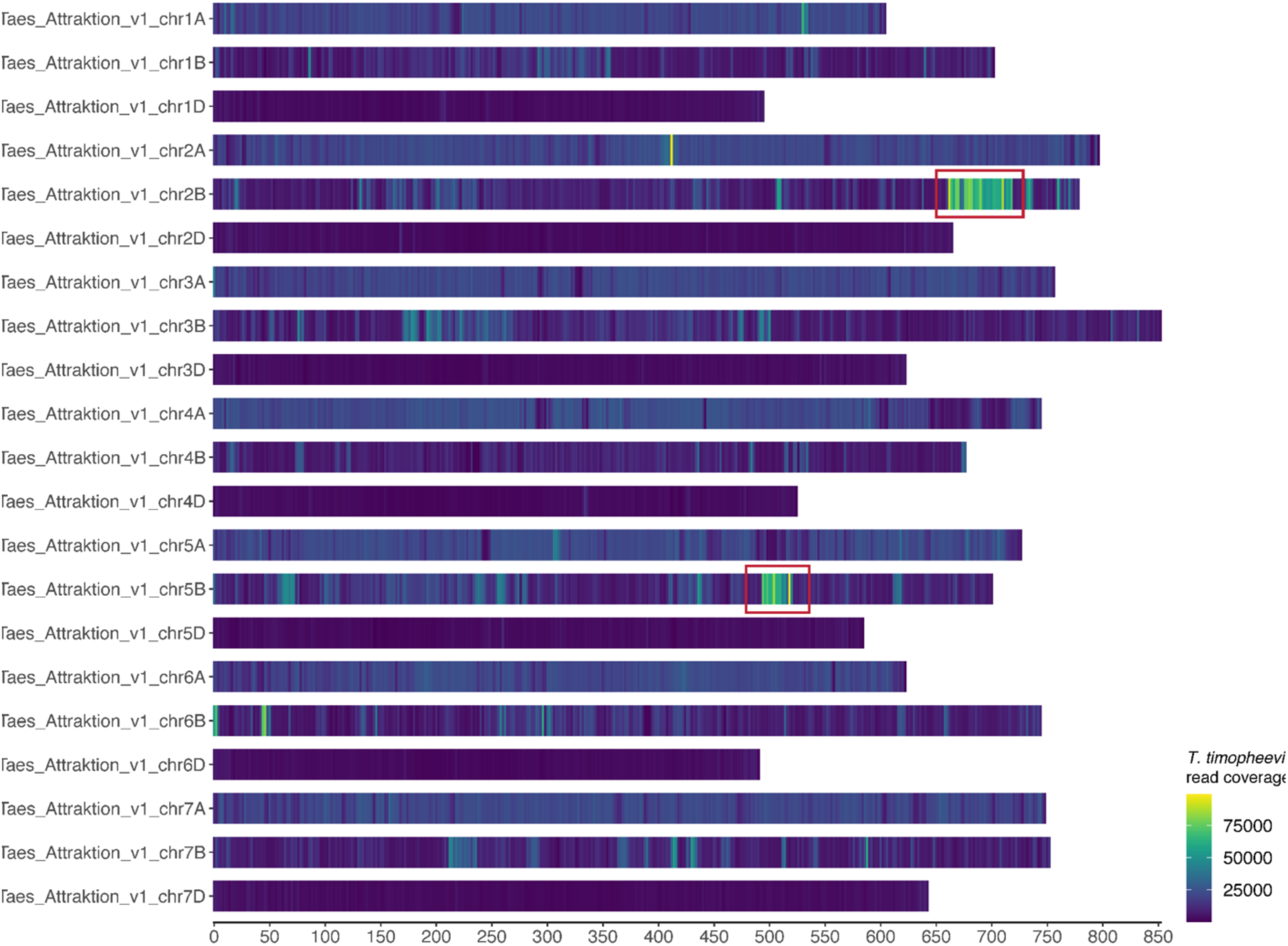
Mapping of wild *T. timopheevi* onto the Attraktion genome assembly. Heatmap showing the read coverage of *T. timopheevi* WGS reads mapped on the genome assembly of Attraktion. The known *T. timopheevi* introgression on Chr. 2B and the introgression on Chr. 5B are highlighted by a red rectangle. Values were calculated in 2 Mb bins.

**Supp. Fig. 7.**
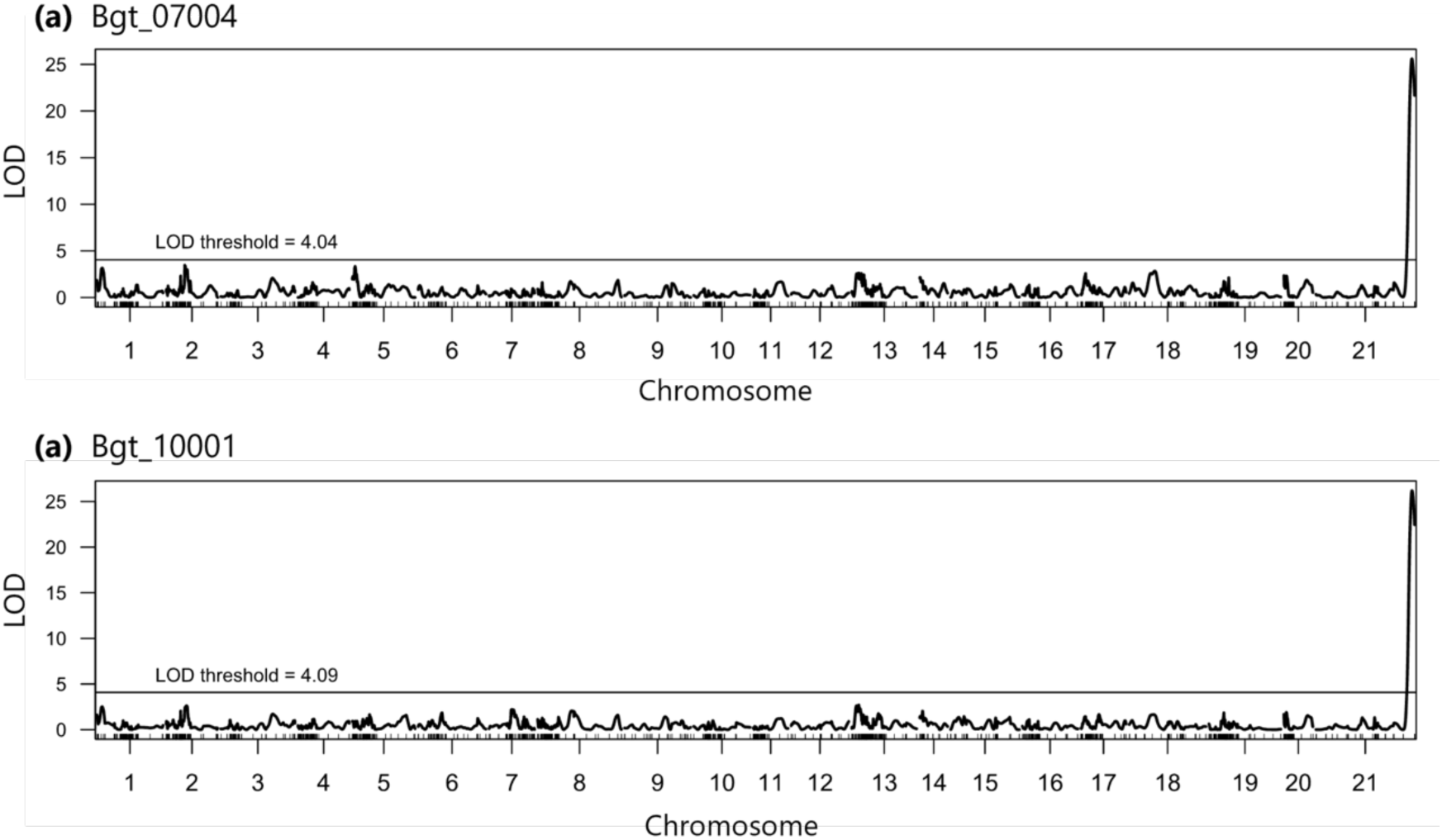
Identification of a powdery mildew resistance locus in chromosome 7D (chromosome 21 in the figure). The powdery mildew isolates **(a)** Bgt_07004 and **(b)** Bgt_10001 reveal the same QTL. The LOD thresholds are indicated in the graphs and were calculated with the *qtl* package in R with 500 permutations.

**Supp. Fig. 8.**
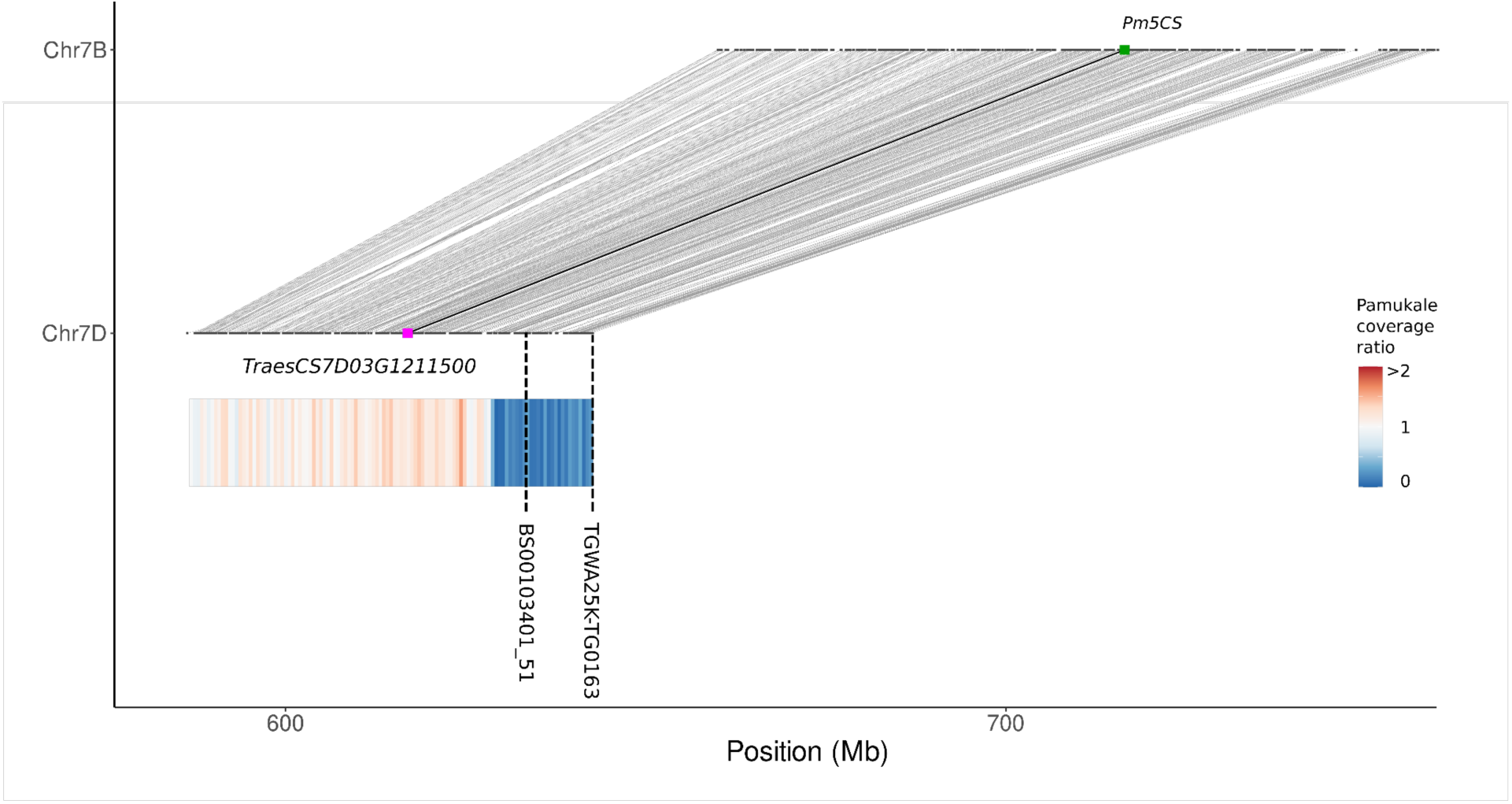
Synteny analysis between the ends of Chinese Spring chromosomes 7B and 7D. Synteny was determined based on blastn queries of the respective gene coding sequences against each other. Pm5CS and its 7D homoeolog are indicated with a green and a pink dot respectively. The position of two SNP markers associated with the *PmPam* QTL is indicated by dashed lines.

**Supp. Fig. 9.**
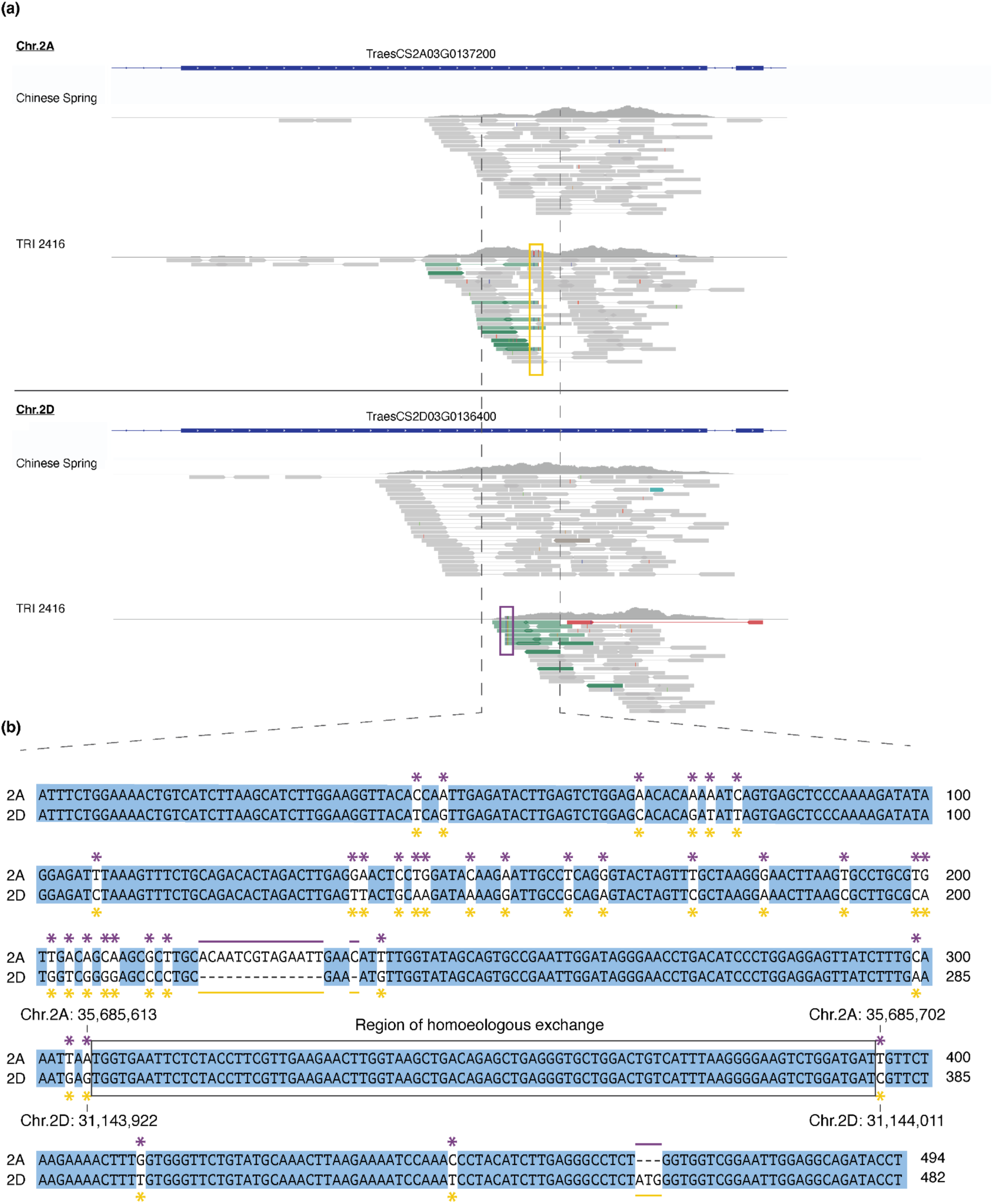
Identification of inter-homoeolog recombination break-point. **(a)** Visualization of mapped reads from Chinese Spring and TRI 2416 exported from IGV. Mate-pairs going across the break-point (one mate mapping to Chr.2A the other to Chr.2D) were coloured in green. Reads that show both 2A- and 2D-specific SNPs in the same read (i.e. chimeric reads) and framed by coloured rectangles. **(b)** Alignment of Chr.2A and Chr.2D in the region surrounding the suspected break-point 2A- or 2D-specific SNPs are indicated by asterisk and specific InDels are indicated by coloured lines.

**Supp. Fig. 10.**
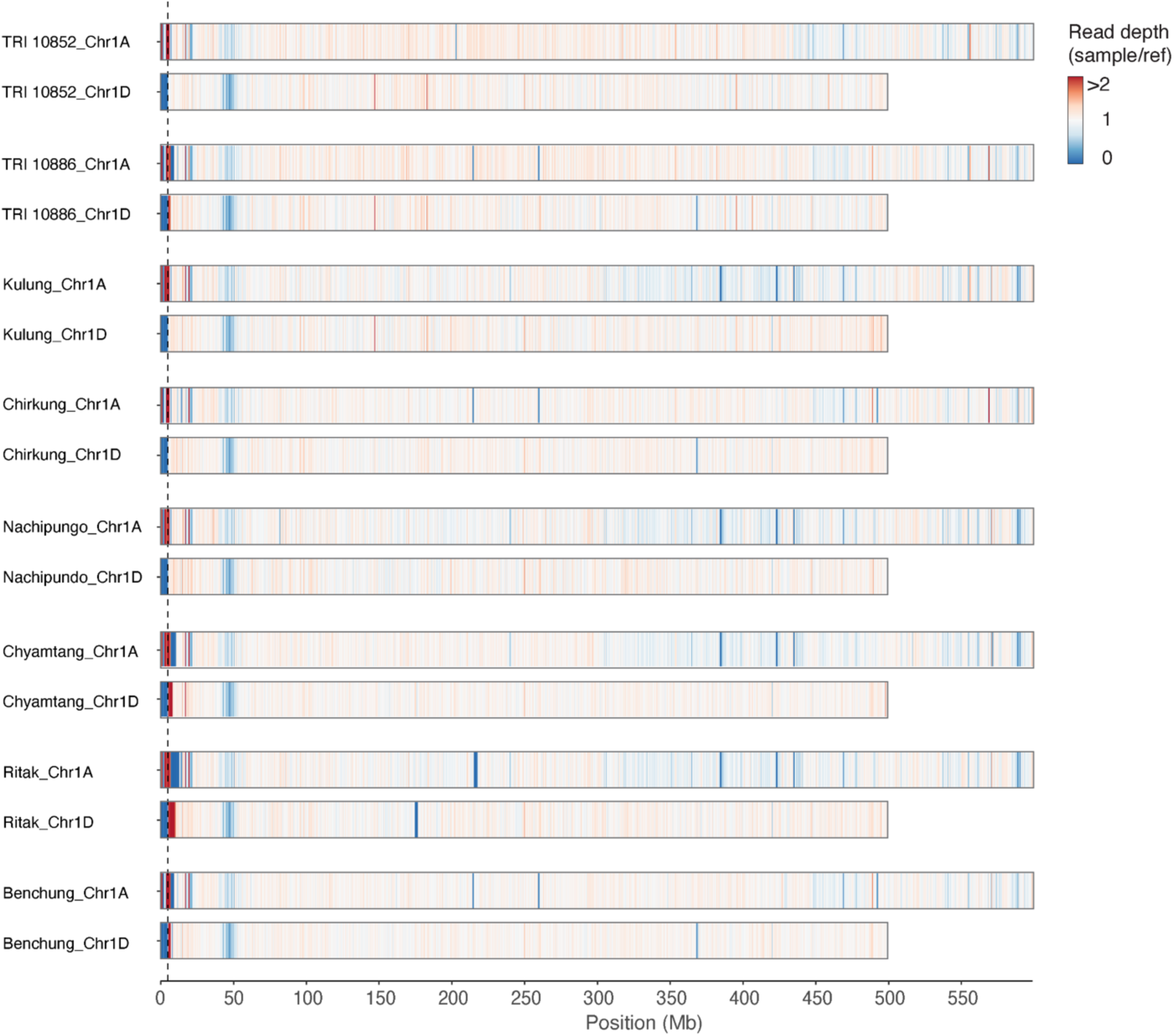
Suspected inter-homoeolog recombination event common to eight Nepalese accessions. Exome capture reads were mapped to the genome of Chinese Spring and coverage ratio was calculated in 500 kb bins. The 8 shown accessions all share one inter-homoeolog recombination event between chromosome 1A and 1D. The location of the breakpoint in chromosome 1D is indicated by a dashed line.

**Supp. Fig. 11.**
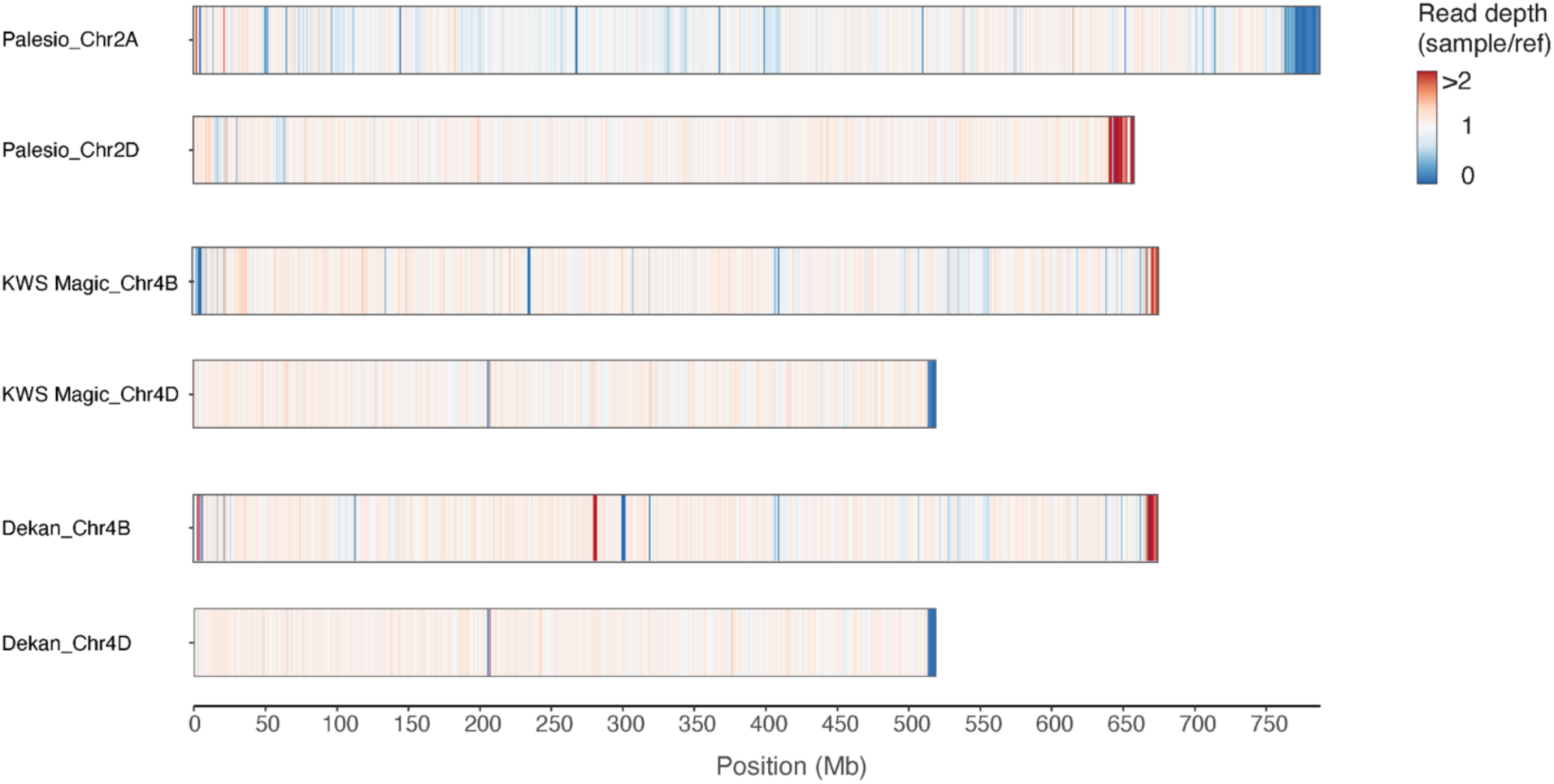
Suspected inter-homoeolog recombination events in current varieties. Exome capture reads were mapped to the genome of Chinese Spring and coverage ratio was calculated in 500 kb bins.

